# Thrombospondin-1 promotes fibro-adipogenic stromal expansion and contractile dysfunction of the diaphragm in obesity

**DOI:** 10.1101/2023.08.17.553733

**Authors:** Eric D. Buras, Moon-Sook Woo, Romil Kaul Verma, Sri Harshita Kondisetti, Carol S. Davis, Dennis R. Claflin, Kimber Converso Baran, Daniel E. Michele, Susan V. Brooks, Tae-Hwa Chun

## Abstract

Pulmonary disorders impact 40-80% of individuals with obesity. Respiratory muscle dysfunction is linked to these conditions; however, its pathophysiology remains largely undefined. Mice subjected to diet-induced obesity (DIO) develop diaphragmatic weakness. Increased intra-diaphragmatic adiposity and extracellular matrix (ECM) content correlate with reductions in contractile force. Thrombospondin-1 (THBS1) is an obesity-associated matricellular protein linked with muscular damage in genetic myopathies. THBS1 induces proliferation of fibro-adipogenic progenitors (FAPs)—mesenchymal cells that differentiate into adipocytes and fibroblasts. We hypothesized that THBS1 drives FAP-mediated diaphragm remodeling and contractile dysfunction in DIO. We tested this by comparing effects of dietary challenge on diaphragms of wild-type (WT) and *Thbs1* knockout (*Thbs1^-/-^*) mice. Bulk and single-cell transcriptomics demonstrated DIO-induced stromal expansion in WT diaphragms. Diaphragm FAPs displayed upregulation of ECM and TGFβ-related expression signatures, and augmentation of a *Thy1*-expressing sub-population previously linked to type 2 diabetes. Despite similar weight gain, *Thbs1^-/-^* mice were protected from these transcriptomic changes, and from obesity-induced increases in diaphragm adiposity and ECM deposition. Unlike WT controls, *Thbs1^-/-^*diaphragms maintained normal contractile force and motion after DIO challenge. These findings establish THBS1 as a necessary mediator of diaphragm stromal remodeling and contractile dysfunction in overnutrition, and potential therapeutic target in obesity-associated respiratory dysfunction.

## INTRODUCTION

Obesity affects over 40% of Americans [1], predisposing them to respiratory disorders that include dyspnea on exertion (DOE) and obesity hypoventilation syndrome (OHS). DOE impacts 30-80% of people with obesity, while OHS prevalence ranges from 10% in individuals with body mass index (BMI) of 30-35 kg/m^2^ to more than 50% in those with BMI >50 kg/m^2^ [2-11]. DOE reduces the quality of life and impairs exercise tolerance [12-14], while OHS confers 5-year heart failure and mortality rates twice those of demographically matched controls [15]. Exercise therapy may mitigate DOE; however, OHS treatment remains medically challenging: Aside from significant weight loss, chronic non-invasive positive pressure ventilation (NIPPV) is its therapeutic mainstay; and permanent tracheostomy is required in severe cases [16].

Clinical studies implicate dysfunction of the respiratory muscles—most notably the diaphragm—as a driver of obesity-associated respiratory impairment [17-20]. While its underlying pathophysiology remains unclear, correlations between disordered breathing and increased limb muscle adiposity suggest diaphragm muscle quality may be compromised in people with elevated BMI [21-23]. To this end, an autopsy study identified large adipocyte inclusions in the diaphragms of individuals with OHS [24]. Intriguingly, patients with obesity also have significantly higher mortality rates following COVID-19 infection [25]—a condition recently shown to promote diaphragm fibrosis [26].

We previously applied a mouse model of long-term diet-induced obesity (DIO) to define the relationship between anatomic remodeling and physiologic dysfunction of the diaphragm. In mice subjected to 6-month high fat diet (HFD) feeding, we found that diaphragm contractile strength declines and inversely correlates with intramuscular adipocyte number and polymerized collagen content [27]. In HFD-fed mice, PDGFRα-expressing fibro-adipogenic progenitors (FAPs) are key contributors to intra-diaphragmatic adipocyte accumulation and extracellular matrix (ECM) deposition [27].

FAPs are mesenchymal stem cells that reside within skeletal muscle and give rise to intramuscular adipocytes and fibroblasts [28-30]. Important regulators of muscle development and maintenance, FAPs orchestrate muscle stem cell (MuSC) activation essential for tissue growth and repair [31-33]. Conversely, in mouse models of muscular dystrophy [34, 35] and severe injury [32, 36], disordered FAP dynamics contribute to pathological adiposity and fibrosis that accompany and potentiate contractile dysfunction [37]. Recent analyses using single-cell RNA-sequencing (scRNA-seq) demonstrate the presence of distinct FAP populations with pro-remodeling and pro-adipogenic potentials [38]. Furthermore, a specific FAP subset marked by *THY1* (CD90) expression is associated with fibro-fatty degeneration in quadriceps muscles of individuals with type 2 diabetes [39]. Despite these recent advances, the molecular mechanisms underlying FAP dysregulation in obesity remain largely unknown.

Thrombospondin-1 (THBS1 or TSP-1) is a matricellular protein present in tissues and circulation. In humans, serum THBS1 levels increase with body mass index (BMI) and are associated with adipose inflammation, insulin resistance, and diabetes [40-43]. In mice, *Thbs1* ablation protects against high-fat diet (HFD)-induced adipose fibrosis while reducing collagen deposition in limb muscles [44]. *In vitro*, THBS1 induces the proliferation of bone marrow-derived mesenchymal cells by potentiating known FAP growth factor TGFβ [45]. Furthermore, clinical and rodent studies link local and circulating THBS1 to fibrotic and degenerative changes in heritable myopathies [46, 47]. We previously demonstrated that THBS1 circulates at high levels in mice subjected to DIO protocols [27, 44] and induces the proliferation of diaphragm-derived FAPs [27]. Therein, THBS1 is a putative regulator of FAP phenotype and consequent diaphragm remodeling in obesity.

Using the *Thbs1*-null state as an interrogating probe, we aimed to identify FAP subtypes involved in obesity-induced diaphragm remodeling; and determine whether *Thbs1* ablation could ameliorate attendant contractile impairment. Our findings indicate that *Thbs1* plays a crucial role in obesity-induced upregulation of TGFβ signaling in diaphragm FAPs while promoting the expansion of a THY1^+^ FAP subtype. *Thbs1* knockout mice are protected from obesity-induced fibro-adipogenic diaphragm remodeling and respiratory dysfunction.

## RESULTS

### Diaphragm FAP transcriptomic profile and sub-populations change in response to diet-induced obesity

Single-cell RNA sequencing has shown FAPs to be a heterogeneous cell type with functionally relevant sub-populations that expand and contract in response to denervation, muscular dystrophy, and injury [48-50]. Transcriptomic changes underlying obesity-associated diaphragm remodeling, on the other hand, remain undefined. To analyze these, we applied scRNA-seq to mononuclear isolates from costal diaphragms (excluding central tendon and rib attachments) of male C57Bl/6J mice subjected to 6-month control diet (CD) (n= 2) or HFD (n= 2) feeding. We computationally aggregated 10X Genomics Cell Ranger data to produce a dataset comprising 3.4x10^8^ reads over 7,906 cells. These resolved into populations including muscle stem cells (MuSCs— expressing *Myf5*, *Pax7,* and *Cdh15*), endothelial cells (expressing *Flt1*, *Pecam1,* and *Ptprb1*), lymphatic cells (expressing *Clca3a1*, *Ccl21a,* and *Mmrn1*), Schwann cells (expressing *Mpz*, *Ncma*p, and *Kcna1*) and macrophages (Mφs—expressing *Itgam*, *Cd68,* and *Lyz2*); as well as a heterogeneous leukocyte cluster containing lymphocytes and eosinophils (Figure 1A, S1A). FAPs—identifiable based on their expression of *Pdgfra*, *Pdgfrb*, *Dcn*, and *Osr1* [29, 51]—were the dominant costal diaphragm cell type, accounting for >50% of sequenced events (Fig 1A-B, S1A). A small mesothelial population (with unique enrichment of *Msln*, *Lrn4,* and *Upk3b*) also expressed *Osr1* and *Dcn* but lacked *Pdgfra* and *Pdgfrb* (Figure 1B, S1A).

**Figure 1:**
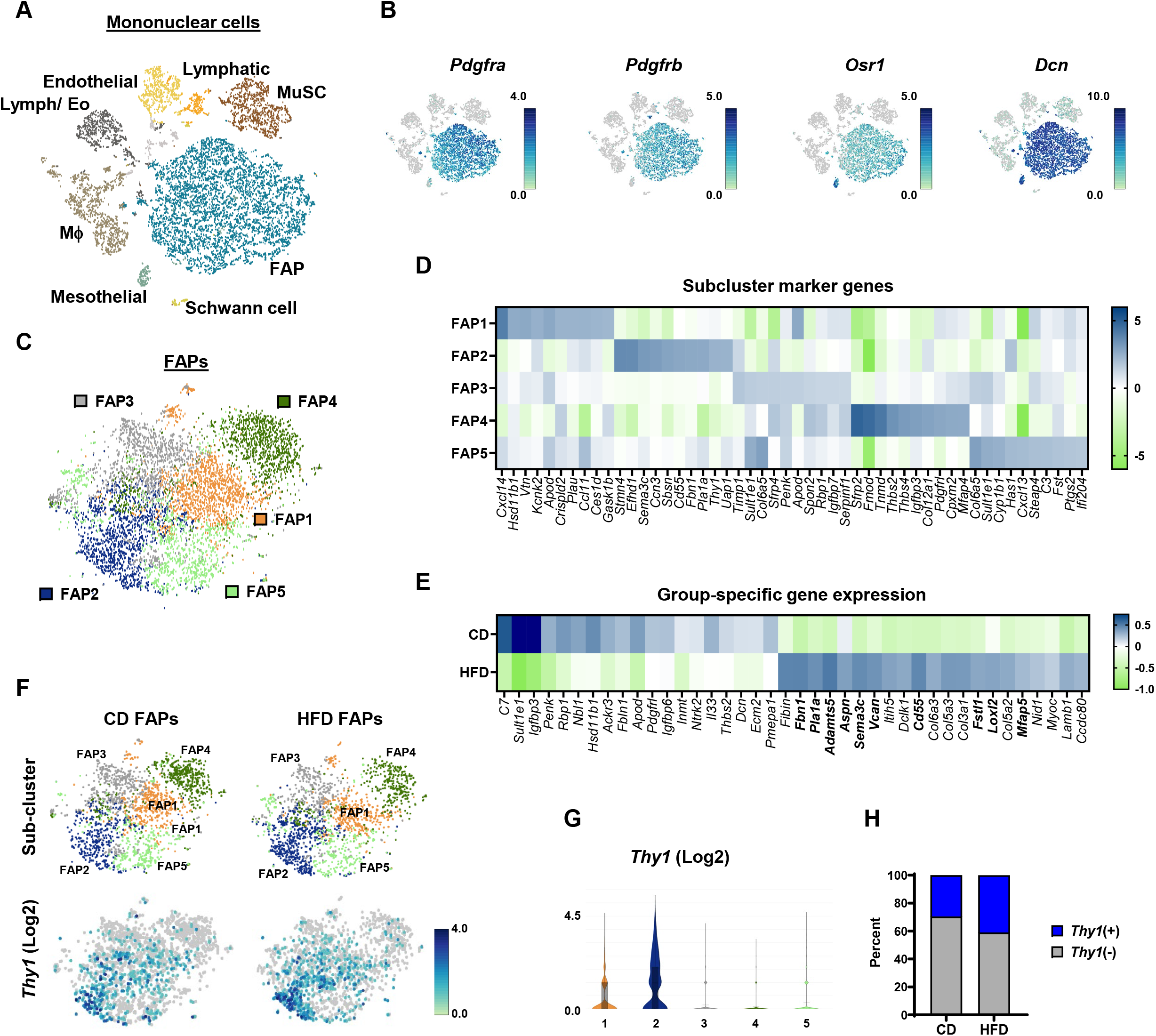
The diaphragm FAP population is heterogeneous and altered by obesity. **(A)** tSNE plot: Mononuclear cells from pooled diaphragms of 6-month high-fat diet (HFD)-fed and age-matched control diet (CD)-fed male C57Bl/6J mice n= 2 mice per group. **(B)** tSNE plots: FAP marker genes. **(C)** tSNE plots: FAP sub-populations from pooled CD and HFD samples. **(D)** Heat map showing transcripts enriched in FAP sub-clusters. **(E)** Heat map showing genes enriched in total FAP populations from CD and HFD samples. Bolded gene names are enriched in the FAP2 sub-cluster. **(F)** tSNE and violin plots showing FAP sub-clusters and *Thy1* expression in CD and HFD samples. **(G)** Violin plots indicating cluster-specific *Thy1* expression. **(H)** Percentage of *Thy1*-expressing among FAPs from CD and HFD samples.

To interrogate their heterogeneity, we re-clustered FAPs into five sub-populations (Figure 1C-D). The largest of these, FAP1, was enriched in transcripts encoding *G0s2*, *Hsd11b1*, *Vtn,* and *Ccl11* (Figure 1D, S1B-C). This expression signature closely resembles the “adipocyte progenitors” defined by Hepler et al. on scRNA-seq analysis of PDGFRβ^+^ cells from mouse adipose tissue [52]. Consistent with this, cells expressing *Mme*—a newly defined marker of adipogenic FAPs [53]—were almost exclusively restricted to FAP1 (Figure S1C). Few FAPs in any sub-cluster expressed *Cebpa, Pparg,* or *Adipoq* (S1D), indicating that the diaphragm FAP pool contains numerous adipocyte progenitors but few committed preadipocytes.

Cells within FAP2 expressed established FAP markers (*Cd55* and *Fbn1*), as well as *Limch*, *Efhd1*, *Smn4*, *Has2*, *Cmah* and *Dact2* (Figure 1D, S2A-C). The latter signature is consistent with the non-adipogenic, ECM-depositing “fibroinflammatory progenitor” population previously described in adipose tissue [52]. In agreement with this, FAP2 cells were enriched in the transcript encoding ECM protein fibronectin (*Fn1*) (FigS2C). Moreover, the FAP2 profile broadly overlapped with that of the FBN1^+^ FAP population identified by Rubenstein et al. in mouse and human muscles [51]. Interestingly, FAP2 cells also demonstrated enhanced (though not exclusive) expression of genes encoding DPP4 (*Dpp4*) (Fig S2B), an adipocyte stem cell marker [54], and endosialin (*Cd248*) (Fig S2D), an obesity-associated glycoprotein that confers adipocyte insulin resistance [55].

Cells within FAP3 highly expressed *Timp1*, a key autocrine regulator of mesenchymal stem cell identity [56]. FAP3 cells were further enriched in transcripts expressed within the FAP5 sub-population, such as that encoding sulfotransferase *Sult1e1* (Figure 1D, S3A-B). Cells within FAP4 had a more specific signature, expressing transcripts encoding secreted regulators of MuSC differentiation such as the small interstitial leucine-rich proteoglycan fibromodulin (*Fmod*) and soluble Wnt signaling modulator SFRP2 (*Sfrp2*) [57, 58]. While FAP4 uniquely contained cells that expressed tenomodulin (*Tnmd*), none within the population expressed the tenocyte marker scleraxis (*Scx*) [59, 60] (Figure 1D, S3C-D).

To assess the impact of dietary modification on the diaphragm FAP population, we analyzed differential gene expression between FAPs from diaphragms of CD and HFD-fed mice. Consistent with our previous work— which showed some diaphragm FAPs to assume a pro-fibrotic phenotype in the setting of DIO [27]—FAPs from HFD samples exhibited enhanced expression of genes encoding ECM species, such as *Col3a1*, *Col5a2*, *Col5a3,* and *Col6a3 (*Figure 1E). In addition, HFD FAPs were enriched in FAP2 transcripts, represented by *Fbn1*, *Pla1a1*, *Sema3c*, *Cd55*, *Mfap5* and *Thy1* (Figure 1E, S2A, C). The latter was particularly intriguing, given a recent report describing an increase in THY1^+^ FAPs in the vastus lateralis muscle of obese individuals with type 2 diabetes. In that study, THY1^+^ FAPs exhibited increased proliferation and fibrogenic differentiation *in vitro*; and were associated with fibro-fatty degeneration in the muscle [39]. In our dataset, *Thy1* expression was largely restricted to FAP2 (Figure 1F-G), while *Thy1*-expressing cells likewise expressed FAP2 marker genes (Figure S4A-B). Notably, *Thy1*-expressing FAPs were more numerous in samples from HFD-fed mice than CD-fed mice (Fig 1H). Taken together, these data demonstrate that diaphragm FAPs resolve into distinct sub-populations, suggesting compartmentalization of adipogenic, ECM-deposition, and regulatory functions. HFD feeding promotes a pro-fibrotic transcriptomic signature and enrichment in the FAP2 subpopulation, with a corresponding increase in *Thy1*-expressing cells. These findings corroborate our earlier demonstration of increased fibrogenic FAPs within the obese diaphragm [27] while revealing similarities between the diaphragms of DIO mice and limb muscles of obese human subjects with type 2 diabetes [39].

### THBS1 underlies quantitative and qualitative changes in diaphragm FAPs during the DIO challenge

We next sought to identify molecular regulators of DIO-induced FAP profile changes; and focused on the obesity-associated stromal cell growth factor THBS1 [27, 43-45]. To ascertain the impact of THBS1 on FAP biology *in vitro*, we isolated FAPs with fluorescence-activated cell sorting (FACS) using an established surface marker profile (Sca-1^+^, CD31^-^, CD45^-^, integrin α7^-^) [28, 29]. We then applied THBS1 at a concentration observed in the plasma of human subjects with obesity [43]. THBS1 administration induced FAP proliferation, as indicated by the increased number of Ki67-labeled cells (Figure 2A). Furthermore, THBS1-treated cells demonstrated enhanced extracellular deposition of the FAP2-associated ECM protein fibronectin (Figure 2B).

**Figure 2:**
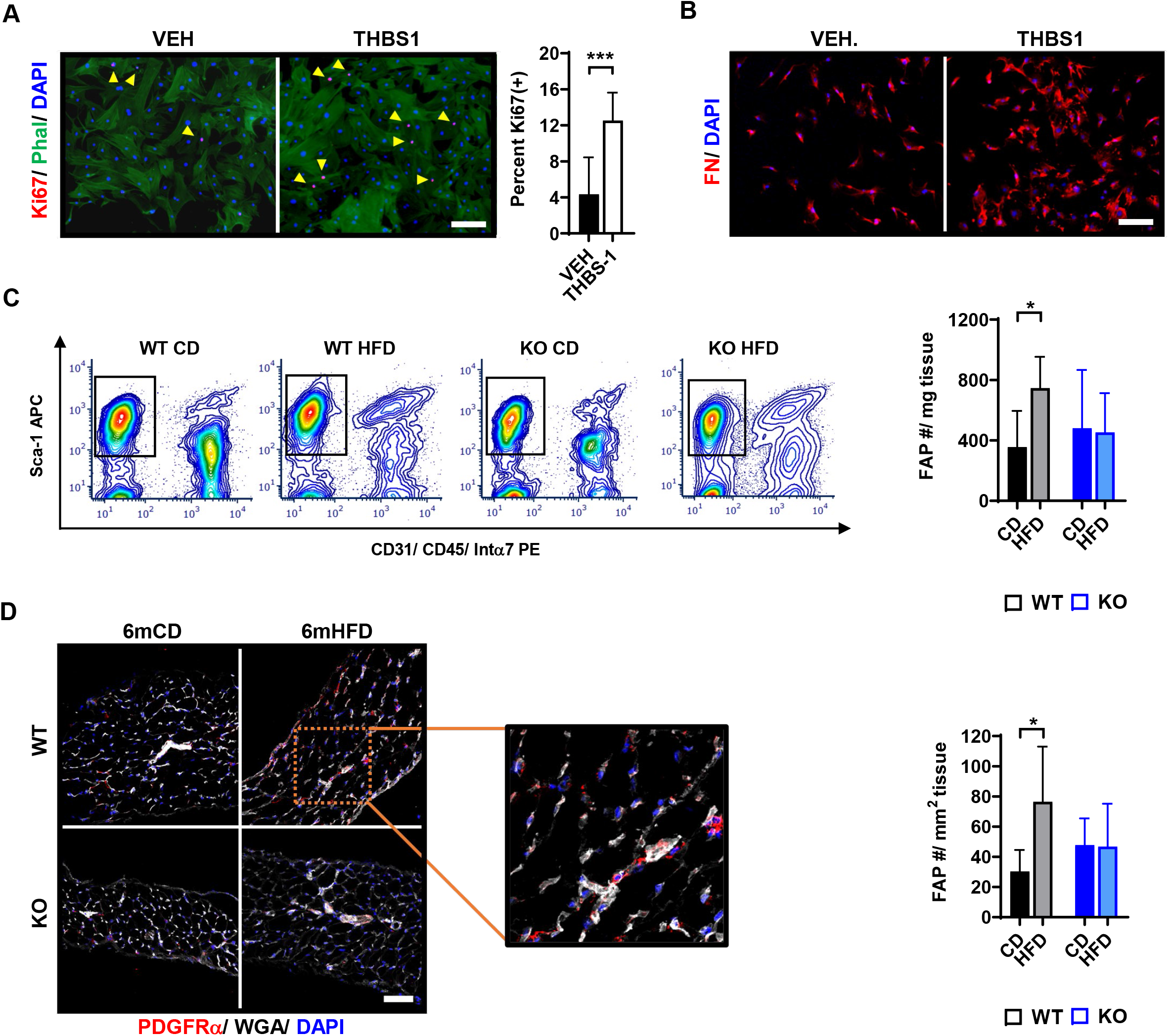
HFD-feeding causes THBS1-dependent FAP population expansion. **(A)** Primary FACS-isolated FAPs treated with THBS1 (5μg/mL) or DMEM vehicle (VEH) and subjected to Ki67 immunocytochemistry with phalloidin (PHAL) counterstain. Scale 50 μm. The bar graph indicates percent Ki67^+^ cells. n= 2 unique experiments per group with 3-4 replicates per experiment. **(B)** Primary FAPs treated as indicated in (A) and subjected to fibronectin (FN) immunocytochemistry. Representative images from 2 unique experiments with 3-4 replicates per experiment. Scale 50 μm. **(C)** Analysis of FAPs from costal diaphragm tissue of wild type (WT) and *Thbs1^-/-^* (KO) mice fed a control diet (CD) or high-fat diet (HFD) for 6 months. Left panels show representative flow cytometry plots—FAPs are positive for Sca-1 and negative for CD31, CD45, and integrin α7 (Intα7). Right panel shows bar graph quantifying FAPs per mg tissue. Each sample contains 2 whole costal diaphragms. n= 5-8 samples (10-16 mice) per group. **(D)** Immunohistochemistry for PDGFRα, with wheat germ agglutin (WGA) counterstain, in diaphragm samples from WT and KO mice fed CD or HFD for 6 months. Bar graph indicates the quantification of PDGFRα^+^ cells/ mm^2^ tissue cross-sectional area (CSA). n= 5-8 mice per group. Error bars indicate SDM. *p<0.05, ***p<0.001.

Given the high level of circulating THBS1 in HFD-fed mice [27], we hypothesized that THBS1 contributed to DIO-associated FAP changes in the diaphragm. To test this, we subjected total *Thbs1* knockout mice (KO mice) [44] (Figure S5A-B) and wild-type (WT) controls to a 6-month HFD-feeding protocol, then evaluated diaphragm FAPs by flow cytometry, immunohistochemistry (IHC) and scRNA-seq.

As previously reported [42], KO mice displayed slightly reduced linear body growth during adulthood versus WT animals (Figure S5C). Nonetheless, they gained weight with HFD feeding (Figure S5D-E) such that their body composition was equivalent to that of HFD-fed WT mice at the end of 6-month dietary manipulation (Figure S5F). Additionally, both WT and KO mice fed HFD showed similarly impaired glucose tolerance compared to age-matched WT mice fed CD (Figure S5G). WT and KO animals also demonstrated comparable DIO-induced increases in liver, perigonadal adipose, and inguinal adipose weights (Figure S6A-C). Finally, weights of several limb muscles did not significantly differ between groups regardless of diet (Figure S6D-H).

Using flow cytometry, we compared mononuclear isolates (two whole costal diaphragms, excluding central tendon and rib attachment, per sample) from WT and KO mice fed either CD or HFD for six months (n= 5-8 samples per group); and quantified FAPs per mg tissue. In WT mice, FAP quantity increased significantly with HFD feeding. On the contrary, KO mice had an equivalent FAP number per mg tissue, regardless of dietary condition (Figure 2C). We corroborated these findings by performing IHC for FAP marker PDGFRα on frozen costal diaphragm samples from mice in the same groups (n= 5-8 animals per group)—observing that number of FAPs per mm^2^ cross-sectional area increased with HFD-feeding in WT but not KO mice (Figure 2D). We then asked whether KO mice were also protected from DIO-induced changes in FAP transcriptomic signature. To assess this, we integrated samples from KO animals fed HFD for 6 months into our scRNA-seq analysis framework. In diaphragm mononuclear isolates from WT CD (n= 2), WT HFD (n= 2) and KO HFD (n= 2) mice (5.2 x 10^8^ reads over 12,275 cells), we focused our attention on FAPs. Velocity analysis of these data demonstrated DIO to induce differentiation from *Timp1*-expressing FAP3 progenitors toward FAP2. In WT mice, this produced a quantitative increase in the FAP2 population, and a commensurate increase in *Thy1*-expressing cells (Fig 3A). In contrast, HFD-fed KO mice were protected from these population shifts and maintained FAP2 and *Thy1*-expressing cell content similar to that of WT mice fed CD (Figure 3A). Consistent with this, total FAP samples from WT HFD mice showed enrichment of marker genes for both FAP2 (Figure 3B) and *Thy1*-expressing cells (Figure S7A) compared to the other groups.

**Figure 3:**
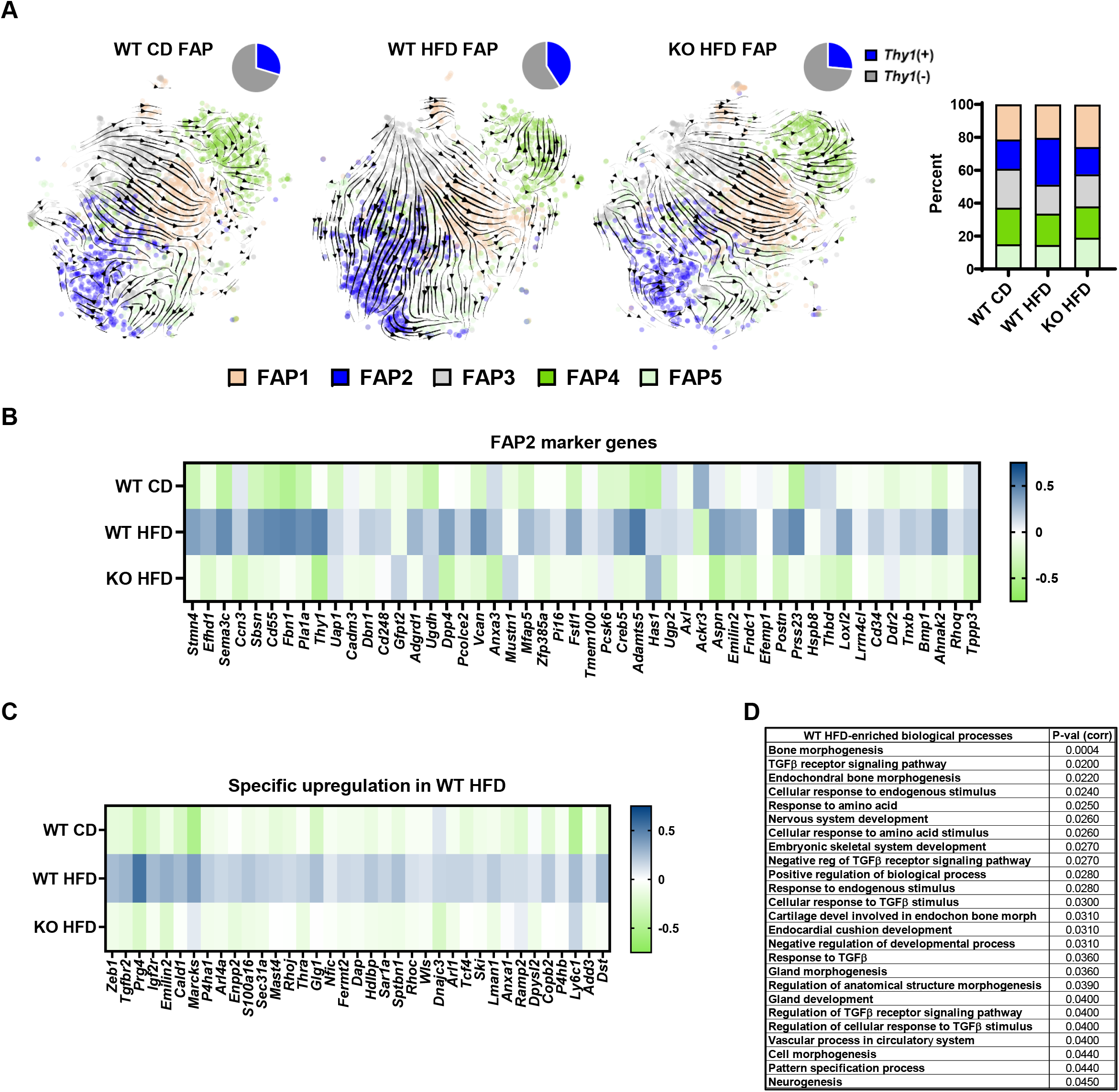
*Thbs1* is required for obesity-induced FAP sub-population shifts. **(A)** Velocity plots demonstrating temporal relationships between FAP sub-populations in wild-type (WT) mice fed CD or HFD for 6 months; and *Thbs1^-/-^* (KO) mice fed HFD for 6 months. Arrow directions represent the trajectory of differentiation between sub-populations. Thickness of arrow indicates a rate of change. Pie charts indicate percentage of *Thy1*-expressing cells in each group. Stacked bar graph shows proportion of individual FAP sub-populations in each group. n= 2 mice per group. **(B)** Heat map indicating the expression of FAP2 marker genes in each group. **(C)** Heat map showing genes specifically enriched in the WT HFD group. **(D)** Gene ontology terms (Biological Processes) specifically enriched in WT HFD FAPs, with corrected p-values.

We next used iPathwayGuide to further examine genes and biological pathways enriched in each group (Figure S7B). Several ECM-related genes—including *Adamts5*, *Sema3c*, *Itih5*, *Cd55*, *Fbn1*, *Col3a1*, *Col4a1*, *Col4a2* and *Col6a3*—were upregulated in both HFD-fed WT and HFD-fed KO FAPs compared to those of WT mice fed CD—however, their expression levels were highest in WT HFD samples (Figure S7C). Genes with specific upregulation in the WT HFD group included those encoding proteins involved in TGF-β signaling and myofibroblastic transition (e.g. *Zeb1*, *Tgfbr2* and *Prg4*) [61-63] (Figure 3C, S7D-E). In agreement with this, numerous TGF-β-associated pathways were specifically enriched in WT-HFD (Figure 3D, S8A) but not in WT CD or KO HFD samples (Figure S8A-E).

Comparison of TGFβ receptor expression among FAP subtypes revealed *Tgfbr2* to be enriched in FAP2. On the contrary, other transcripts encoding receptors for TGFβ and PDGF species (i.e. *Tgfbr1*, *Tgfbr3*, *Pdgfra* and *Pdgfrb*) were expressed equally across FAP populations (Figure S9A-B). Interestingly, the gene encoding CD47—a cell surface THBS1 receptor that also acts as a “don’t eat me signal” capable of inhibiting Mφ-mediated phagocytosis of fibroblasts and other cell types [64]—was also enriched in FAP2 cells (Figure S9C). Conversely, the gene encoding established endothelial THBS1 receptor CD36 [65] was negligibly expressed FAPs of all subtypes (Figure S9C).

Together, these data demonstrate that THBS1 is required for DIO-induced expansion of the total FAP pool and its shift toward *Thy1*-expressing, FAP2 cells. Increased TGF-β signaling—known to be elevated in DIO [66] and to be directly activated by THBS1 [45]—parallels this phenotypic switch. Notably, FAP2 cells exhibit unique enrichment of TGFβ receptors and anti-phagocytic molecules.

### Whole tissue transcriptomics highlights reduced stromal gene expression in obese Thbs1^-/-^ mice

To determine whether FAP profile differences between HFD-fed WT and KO mice corresponded with gene expression changes at the tissue level, we performed bulk RNA-seq on whole costal diaphragm samples (excluding central tendon and rib attachment points; n= 3 diaphragms per group) from these animals. This analysis revealed a pronounced difference in the transcript encoding adipocyte marker leptin (*Lep*)—expressed more highly in WT samples (Figure 4A). We performed an integrated analysis to determine whether other differentially expressed transcripts identified by bulk RNA-seq were enriched in any of the specific mononuclear cell types defined by scRNA-seq. This approach showed that numerous genes more highly expressed in HFD-fed WT diaphragm tissue were specifically enriched in FAPs (*Mfap5*, *Dpt*, *Fbn1*, *Cilp*, *Fn1*, *Pmepa1*, *Ctgf*, *Sod3* and *Prg4*) and Mφs (*S100a4*, *C1qb*, *Ccl6*, *Ctss*, *C1qa*, *Pf4*, *Cd44*, *F13a1*, *Cd68*, and *Plin2*) (Figure 4A-B). On the contrary, (with the exception of Schwann cell marker *Mpz*) transcripts enriched in HFD-fed KO samples were not enriched in any mononuclear cell type. In fact, genes relatively overexpressed in the KO diaphragm included well-known myofiber transcripts, like those encoding parvalbumin (*Pvalb*) and alpha-actinin-3 (*Actn3*) (Figure 4A-B). Consistent with this, GSEA demonstrated the HALLMARK epithelial to mesenchymal transition pathway (which contains ECM-related genes) and inflammatory response pathway (which contains Mφ-related genes) to be enriched in WT samples compared to KO samples (Figure 4C). Quantitative PCR (qPCR) analysis substantiated these findings, showing *Thbs1* ablation to reduce levels of the transcripts encoding leptin (*Lep*), PDGFRα (*Pdgfra*), fibronectin (*Fn1*) and collagen 3 (*Col3a1*) in the diaphragm of HFD-fed mice (Fig 4D).

**Figure 4:**
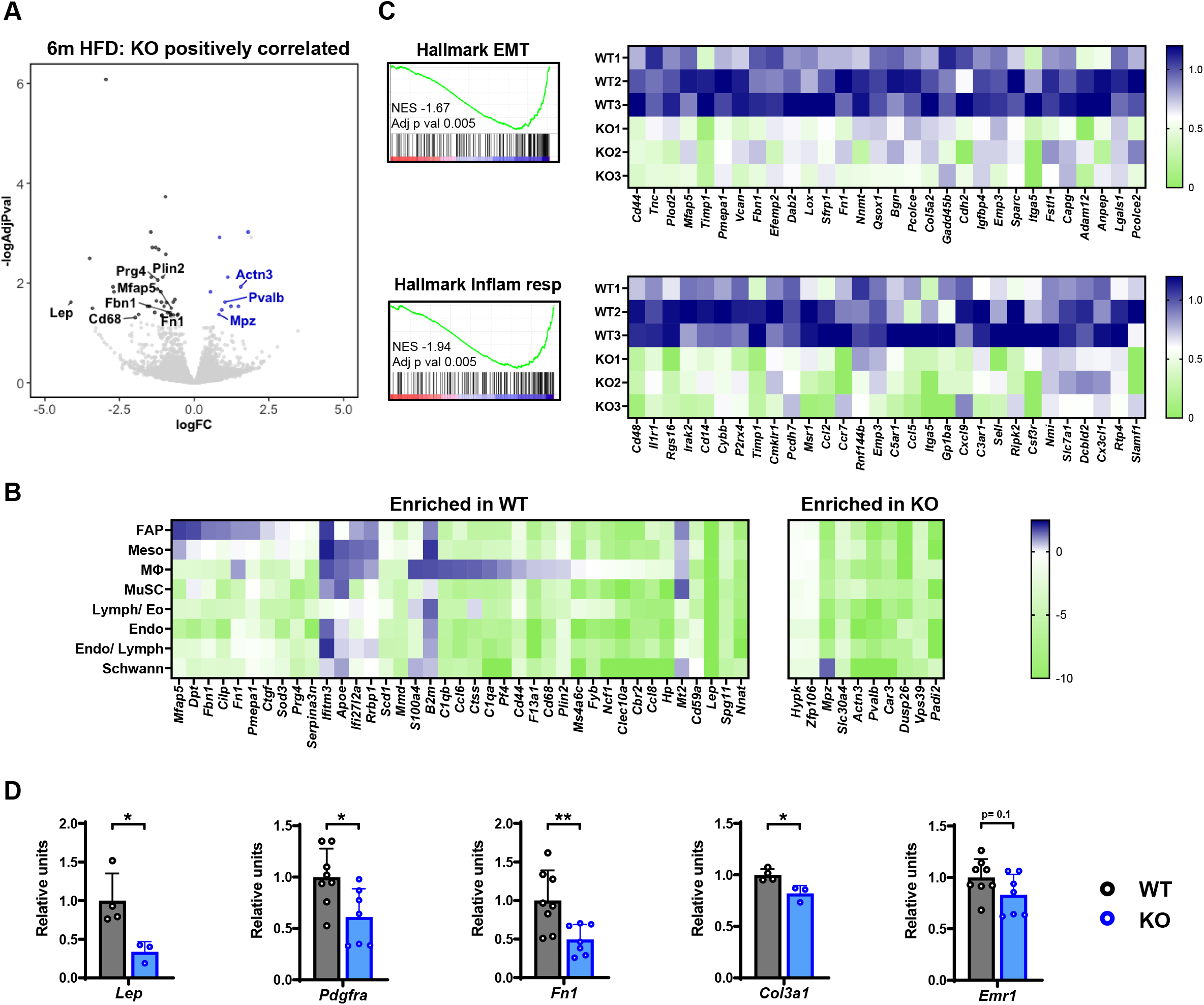
Whole tissue transcriptomics highlights enrichment of stromal genes in wild type versus *Thbs1^-/-^* mice subjected to DIO. **(A)** Volcano plot of whole costal diaphragm RNA-seq demonstrating differentially expressed genes between wild-type (WT) and *Thbs1^-/-^* (KO) mice fed high-fat diet (HFD) for 6 months (6mHFD). n= 3 mice per group. x-axis indicates log fold change (FC). y-axis indicates log adjusted p-value. **(B)** Heat map integrating tissue-level RNA-seq with sc-RNA-seq data. Genes indicated are those enriched in WT and KO mice on bulk RNA sequencing [i.e., the points on the volcano plot in (A)]. Cell types are those identified on scRNA-seq (as shown in Fig 1A). Heat maps show cell-type specific expression as defined on scRNA-seq. **(C)** Enrichment plots demonstrating selected HALLMARK pathways differentially expressed between 6m HFD WT and KO mice. Heat maps show the expression of leading-edge genes in individual samples. **(D)** QPCR analysis of selected genes performed on costal diaphragm tissue of 6mHFD WT and 6mHFD KO mice. Error bars indicate SDM. *p<0.05, **p<0.01.

Expression of *Emr1*, encoding Mφ marker F4/80, also trended down in KO tissue samples; however, the difference did not reach statistical significance (Figure 4D). CD68 immunohistochemistry on frozen costal diaphragm sections further demonstrated HFD feeding to raise Mφ number per mm^2^ tissue cross-sectional area in WT mice. This increase was less pronounced in KO animals (Figure S10A); however, total Mφ number, even in WT HFD-fed mice, was more than four times lower than that of FAPs (Figure 2C). Furthermore, analysis of scRNA-seq data showed lipid-associated *Trem2*-expressing Mφs [67] to exist in equal proportion in samples from WT and KO HFD-fed mice (Figure S10B).

Overall, these data show that, in the obese diaphragm, *Thbs1* ablation reduces expression of stromal genes—particularly those specific to adipocytes, FAPs and Mφs. Of the latter two cell types, FAPs are likely stronger contributors to tissue-level phenotype given their greater numbers. The relative increase in muscle-specific transcripts within the KO diaphragm suggests increased muscle/ stroma ratio in the setting of *Thbs1* ablation.

### Thbs1 ablation protects against diaphragm fibro-adipogenic remodeling

We previously demonstrated that DIO promotes diaphragm tissue remodeling, characterized by increased FAP-derived ECM-depositing cells and intramuscular adipocytes [27]. Given that *Thbs1* ablation ameliorated FAP population expansion and subtype shifts, while reducing expression of adipocyte and ECM-related transcripts, we surmised that KO mice would be protected from the remodeling phenotype.

To assess this, we examined tissue morphology in H/E-stained longitudinal costal diaphragm sections spanning the rib and tendon attachment points (n= 5-8 mice per group, 3-4 non-consecutive sections per animal) (Figure 5A). In WT mice, this analysis demonstrated intramuscular adipocyte inclusions that were particularly prominent in the lateral costal diaphragm and larger in mice subjected to 6-month DIO (Figure 5A). Indeed, intramuscular adipocyte size and number; as well as total tissue cross-sectional area occupied by adipocytes, increased in HFD-fed versus CD-fed animals (Fig 5B). Despite similar weight gain to HFD-fed WT mice, HFD-fed KO mice were largely protected from these obesity-associated changes in intramuscular adiposity (Figure 5A-B).

**Figure 5:**
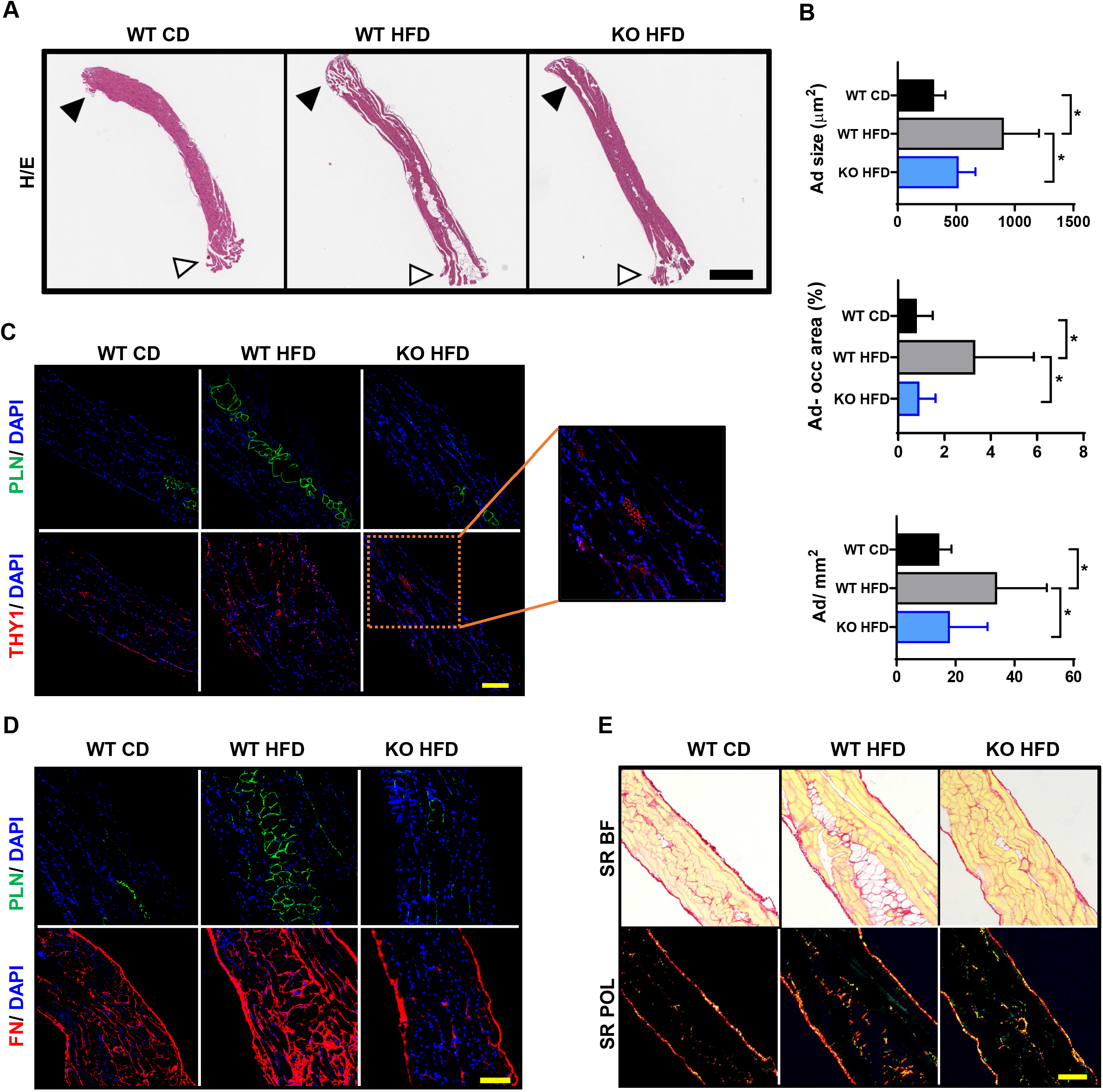
*Thbs1* ablation protects against diaphragm fibro-adipogenic remodeling. **(A)** H/E-stained longitudinal diaphragm sections from wild-type mice fed control diet (WT CD) or HFD (WT HFD) for 6 months; and *Thbs1^-/-^* (KO) mice fed HFD for 6 months (KO HFD). White arrowhead indicates rib attachment point. Black arrowhead indicates central tendon attachment point. Scale 600 μm. Representative samples from 5-6 mice per group. **(B)** Adipocyte size, adipocyte number per mm cross-sectional area, and percent total cross-sectional area occupied by adipocytes in samples described in (A). Values are the average of measurements made on 3 non-consecutive 7 μm-thick sections encompassing the entire rib-to-tendon extent of muscle. n= 5-6 mice per group. **(C)** Immunofluorescent staining of perilipin (PLN) and THY1 on adjacent 7 μm-thick longitudinal sections from animals described above. Representative images from analysis of 5-6 mice per group. Inset indicates THY1 staining of a nerve passing through the sample, representing an internal positive staining control. Scale 200 μm. **(D)** PLN and fibronectin (FN) staining on adjacent 7-μm-thick longitudinal sections from animals described above. Representative images from analysis of 5-6 mice per group. Scale 200 um. **(E)** Picrosirius red (SR) staining of 7 μm-thick longitudinal sections from animals described above. Bright-field (BF) and polarized light (POL) images: polymerized collagens fluoresce red/ yellow under polarized light. Scale 200 μm. Error bars indicate SDM. *p<0.05.

We next sought to understand the geographical relationship between these intramuscular adipose depots and THY1^+^ FAPs. Immunofluorescent staining of adjacent longitudinal sections for adipocyte marker perilipin and THY1 demonstrated collections of THY1^+^ cells close to adipocyte inclusions—their number higher in HFD-fed WT samples versus the other groups (Figure 5C). Moreover, prominent deposition of both fibronectin (Figure 5D) and polymerized collagen (picrosirius red staining) (Figure 5E) surrounded adipose depots. Both increased with HFD-feeding in WT mice, while samples from HFD-fed KO mice resembled those of WT mice fed CD. Notably, we did not observe significant differences in diaphragm thickness (Figure S11A), myofiber cross-sectional area (Figure S11B), or myofiber type (Figure S11C) between WT and KO mice fed HFD.

Together, these histological analyses provide evidence that *Thbs1* ablation protects the diaphragm from DIO-induced fibro-adipogenic remodeling in a manner consistent with the effects predicted from transcriptomic profiles at the cell and tissue level. Moreover, the findings indicate that DIO-induced increases of intramuscular adiposity, ECM deposition and THY1^+^ cells occur within the same anatomic regions of the tissue and depend on THBS1.

### The Thbs1^-/-^ diaphragm preserves its contractile force in the setting of DIO challenge

Given the improved tissue architecture observed in samples from HFD-fed KO versus WT mice, we predicted that *Thbs1* ablation would also protect the diaphragm from obesity-associated mechanical dysfunction. To test this hypothesis, we performed *ex vivo* isometric force testing on diaphragm strips (Figure S11D) isolated from WT and KO mice at baseline (2-months old) and following 6-month CD or HFD feeding (n= 5-7 mice per group, 2 diaphragm samples per mouse). In WT animals, 6-month CD feeding effected no difference in isometric force versus baseline. HFD feeding, on the other hand, caused specific force to decline by approximately 20% (Figure 6A). In KO mice, specific force values remained unchanged from baseline regardless of whether animals received CD or HFD (Figure 6A). In addition, samples from HFD-fed KO mice had significantly higher specific force measurements than those of HFD-fed WT mice. (Figure 6B). Therefore, *Thbs1* ablation protects the diaphragm from obesity-associated contractile force reduction.

**Figure 6:**
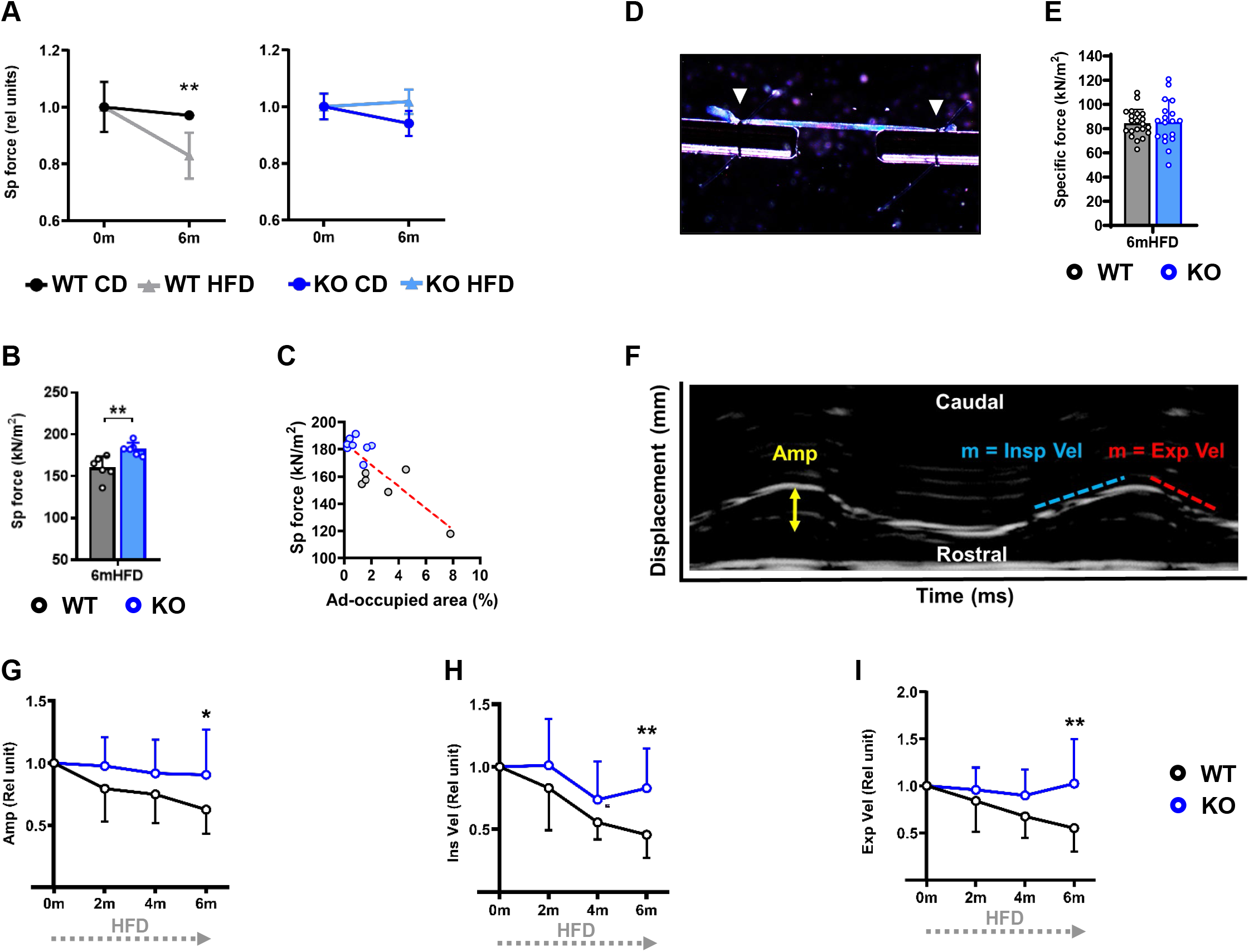
DIO challenge compromises diaphragm force and motion in WT but not *Thbs1^-/-^* mice. **(A)** Isometric specific force of wild-type (WT) and *Thbs1^-/-^* (KO) mice (normalized to baseline) at baseline (0m) and following 6-month (6m) control diet (CD) or high-fat diet (HFD) feeding. n= 4-6 animals per group; 1-2 diaphragm strips per animal averaged. **(B)** Isometric specific force (absolute value) of samples from 6mHFD WT and KO mice. n= 5-6 animals per group; 1-2 diaphragm strips per animal averaged. **(C)** Correlation plot demonstrating the relationship between isometric specific force and percent tissue cross-sectional area occupied by adipocytes in diaphragm strips subjected to isometric force testing. 6mHFD WT and KO mice, 8-9 muscle strips per group. **(D)** Image of single myofiber undergoing isometric force testing. White arrowheads indicate sutures affixing fiber to force transducer-servomotor apparatus. **(E)** Isometric specific force of single myofibers isolated from 6mHFD WT and KO mice (n= 4-5 animals per group; 5 fibers per animal). **(F)** Diaphragm ultrasound M-mode tracing with measured parameters labeled. x-axis represents time, y-axis represents displacement along the rostral-caudal axis. **(G-I)** Diaphragm motion parameters: amplitude (Amp), inspiratory velocity (Ins Vel), expiratory velocity (Exp Vel), normalized to baseline measured at 0m, 2m, 4m and 6m. n= 7-9 animals per group. Error bars indicate SDM. *p<0.05, **p<0.01.

Within diaphragm muscle strips from HFD-fed WT and KO mice subjected to isometric testing, we observed a negative correlation between adipocyte-occupied cross-sectional area and measured specific force (Figure 6C). Given this relationship, we surmised that tissue-level contractile force deficits resulted from altered muscle architecture. An alternative explanation is that THBS1 could directly impair myofiber function in the context of obesity. To test this possibility, we performed isometric force testing on single myofibers isolated from WT and KO mice fed HFD for 6 months (Figure 6D). This analysis found specific force measurements of individual fibers from mice of each group to be statistically indistinguishable. (Figure 6E, S11E). As such, the protective effect of *Thbs1* ablation is not dependent on better sarcomere contractility, but instead may result from undisrupted tissue architecture necessary for coordinated muscle contraction.

### Thbs1 ablation protects mice from obesity-associated deterioration of diaphragm motion

We then asked whether the preservation of normal diaphragm contraction seen in HFD-fed KO mice translated into protection from obesity-associated respiratory dysfunction. To assess this, we subjected WT and KO to a 6-month HFD time course and serially analyzed diaphragm motion with non-invasive ultrasound. M-mode measurements—plotted with time on the x-axis; and diaphragm displacement on the y-axis—enable measurement of diaphragm excursion amplitude, inspiratory velocity, and expiratory velocity (Figure 6F) [27, 68]. In WT mice, each of these parameters progressively declined with HFD feeding duration. On the contrary, in KO mice, all measurements remained stable throughout the time course (Figure 6G-I). Moreover, at the 6-month time point, baseline-normalized amplitude, inspiratory velocity, and expiratory velocity were all significantly higher in KO than WT mice (Figure 6JG-I). In sum, animals lacking *Thbs1* are protected from obesity-associated compromise of diaphragm motion.

## DISCUSSION

Our findings demonstrate that anatomic remodeling and contractile dysfunction of the diaphragm are interrelated, *Thbs1*-dependent obesity complications. In the setting of long-term overnutrition, THBS1 promotes stromal expansion characterized by increased THY1^+^ FAPs, aberrant ECM deposition and elevated intramuscular adiposity. These changes correspond with a decline in tissue-level isometric force generation— independent of single myofiber sarcomere function—that ultimately contributes to reduced diaphragm motion.

### THBS1 underlies obesity-associated alterations in diaphragm FAP profile

Our results define THBS1 as a key regulator of quantitative expansion and qualitative changes in the diaphragm FAP pool during long-term DIO. A circulating matricellular protein, THBS1 is produced by megakaryocytes, platelets, leukocytes, endothelial cells, fibroblasts, and adipocytes [40, 69-71]. In most cell types, THBS1 expression is low at baseline but acutely rises as a component of wound healing and ischemic stress responses [72]. Persistent, maladaptive THBS1 elevation occurs with aging and in the setting of prolonged nutritional stress. For instance, *THBS1* expression increases in adipose depots of obese human subjects and positively correlates with degree of insulin resistance [40]; while plasma THBS1 concentrations are higher in patients with impaired glucose tolerance and HbA1c elevation [43, 73]. Rodent models of metabolic syndrome recapitulate these findings [44], as evidenced by increased circulating THBS1 levels observed in mice subjected to DIO protocols [27, 44].

Deposited in the ECM, THBS1 induces context-specific trophic effects—both proliferation and ECM production—in stromal cells [72]. In cultured bone marrow-derived mesenchymal cells (BMDSCs), THBS1 acts as a potent mitogen [45]. Our *in vitro* data demonstrates an analogous effect on FAPs, as obesity-specific THBS1 concentrations [43] promote their proliferation. Analysis of FAPs from diaphragms of KO mice supports the relevance of this effect *in vivo*: Unlike WT mice, which undergo expansion of the FAP pool with obesity, KO mice maintain similar FAP numbers to baseline when challenged with 6-month DIO. Mechanistically, THBS1 induces proliferation of BMDSCs by activating established FAP mitogens—specifically facilitating the conversion of latent to active TGFβ, and stabilizing PDGF protein [45]. Our transcriptomic analyses highlight TGFβ as a likely driver of *Thbs1*-dependent FAP proliferation in the obese diaphragm, given the enrichment of TGFβ-dependent transcripts and pathways in diaphragm FAPs from HFD-fed WT mice compared to CD-fed WT and HFD-fed KO mice. These data further implicate THBS1 as a driver of augmented TGFβ signaling previously described in muscles of human subjects with metabolic syndrome [74]. Of note, parallel impacts of THBS1 on other trophic factors or on FAP survival [32, 39, 75]—for instance, via reduced Mφ-mediated clearance [64]—might also regulate diaphragm FAP pool size during overnutrition; and therein warrant future exploration.

THBS1 is a crucial mediator of mesenchymal ECM production in numerous tissues [76]—and this process appears operative in diaphragm FAPs. *In vitro*, THBS1 treatment induces deposition of fibronectin, an ECM molecule previously shown to be transcriptionally regulated by TGFβ through c-Jun N-terminal kinase (JNK) signaling [77]. *In vivo*, FAPs isolated from the obese diaphragm assume a fibrogenic signature, with many of the transcripts most enriched in HFD-fed versus CD-fed samples encoding structural or regulatory components of the ECM. THBS1 is required for this shift, since the expression of numerous obesity-induced FAP genes (such as *Fbn1*, *Col3a1* and *Adamts5*) is markedly blunted in samples from HFD-fed KO mice. Moreover, some of the genes most upregulated in WT HFD versus WT CD and KO HFD FAPs include species involved in TGFβ signaling (e.g. *Zeb1*, *Tgfbr2* and *Prg4*).

scRNA-seq-based sub-clustering defined several diaphragm FAP subpopulations; some exhibiting considerable transcriptomic overlap with those described in previous scRNA-seq studies. The most notable was *Thy1*-expressing FAP2—a cell type similar to the fibroinflammatory progenitor described in visceral and subcutaneous adipose tissue and the FBN^+^ FAP subset identified in human and mouse skeletal muscle [51, 52]. In our model, the FAP2 population arose from *Timp1*-expressing FAP3 precursors and expanded with diet-induced obesity. This population shift influenced the transcriptomic profile of the total FAP pool, causing it to become enriched in FAP2 markers. We noted a striking resemblance between FAP2 and the THY1^+^ FAPs shown to increase in vastus lateralis biopsies of individuals with type 2 diabetes. In these human samples, the presence of THY1^+^ cells was associated with fibro-fatty tissue remodeling [39] similar to that observed in diaphragms of DIO mice [27]. THBS1 is critical for the obesity-induced enrichment of FAP2 cells, as subtype profile and velocity plots of FAPs from HFD-fed KO mice were nearly indistinguishable from those of WT mice fed a control diet. Dietary condition and *Thbs1* ablation exerted comparatively little impact on the relative size of the FAP1 subpopulation—an analog of previously-described adipocyte progenitors [52]. The observed FAP profiles therefore align with our previous findings: While DIO induces both increased intra-diaphragmatic adiposity and fibrosis, FAPs isolated from the obese diaphragm do not display enhanced *ex vivo* adipogenesis but do exhibit upregulated ECM deposition. As such, increased intramuscular adipocyte number likely results from increased FAP number [27] and/or THBS1-dependent tissue remodeling creating a microenvironment permissive for adipocyte differentiation and expansion *in vivo*.

On the whole, THBS1 facilitates obesity-associated expansion of the diaphragm FAP pool, inducing a fibrogenic transcriptomic signature typified by TGFβ-dependent gene expression and enrichment of a THY1^+^ subpopulation previously linked to tissue-level remodeling.

### THBS1-induced FAP changes correspond with fibro-adipogenic stromal expansion in the obese diaphragm

During a 6-month DIO time course, the diaphragm undergoes progressive anatomic remodeling characterized by increased intramuscular adiposity and ECM deposition [27]. Here, we show that adipocytes in 6-month DIO mice are not uniformly distributed throughout the costal diaphragm tissue, but instead exist largely in aggregations close to the rib attachment point. ECM distribution is also not homogenous, as densities of polymerized collagens and fibronectin often appear closely interposed with adipocytes. Similarly, THY1^+^ cells preferentially congregate near these intramuscular adipose depots, demonstrating a coupling of their presence and fibro-fatty expansion. *Thbs1* is an essential mediator of these processes, given that KO mice are protected from adipose depot expansion as well as from the associated increase in THY1^+^ FAPs and ECM deposition.

Tissue-level transcriptomic analysis substantiates these histological findings, showing that, compared to the HFD-fed WT diaphragm, the HFD-fed KO diaphragm displays reduced expression of numerous stromal genes, particularly those associated with adipocytes and ECM-depositing FAPs (e.g. *Lep*, *Fn1*, *Fbn1*, *Prg4* and *Mfap5*). Conversely, transcripts more highly expressed in the HFD-fed KO diaphragm include those specific to the myofiber (e.g. *Actn3* and *Pvalb*). This raises questions as to whether increases in myofiber-specific transcripts in the obese KO diaphragm are relative—i.e. occurring because there is less stromal tissue than in the obese WT diaphragm—or represents a protective effect of *Thbs1* ablation on myofiber preservation during overnutrition. Our histological data supports the former possibility, as diaphragm thickness, myofiber size and myofiber type are unchanged in diaphragms of HFD-fed WT and HFD-fed KO mice. Moreover, even compared with diaphragms of CD-fed WT mice, the WT HFD diaphragm does not display pathological hallmarks of atrophy such as centrally nucleated or angular myofibers [27]. That said, we cannot completely rule out slow, progressive, *Thbs1*-dependent myofiber loss coupled with replacement by stromal tissue as a potential driver of obesity-induced diaphragm remodeling and dysfunction.

Regarding THY1^+^ FAPs, our data and prior reports [39, 52] suggest they are predominantly fibrogenic, and therein contribute to the deposition of fibronectin and other ECM species near intramuscular adipose depots. That said, delineating other roles they might play in tissue-level remodeling—e.g. differentiation into adipocytes [53] or secondary promotion of adipose depot expansion through inhibition of myofiber maintenance—are important future directions. Finally, subsequent studies including both male and female mice will be required to determine whether any aspects of obesity-associated diaphragm remodeling or FAP complement are sexually dimorphic [78].

### Thbs1 is necessary for obesity-induced diaphragm and motion contractile deficits

Our testing of isometric specific force in isolated diaphragm strips demonstrates that 6-month DIO markedly impairs contractile function. The process depends on *Thbs1*, as diaphragm samples from KO mice subjected to the same diet maintain equivalent specific force versus baseline and demonstrate significantly greater force than isolates from HFD-fed WT mice. In contrast, measurements of specific force in single myofibers isolated from HFD WT and HFD KO mice exhibited no difference between groups. These data can be interpreted as demonstrating that obesity-induced isometric force deficits result from tissue-level remodeling rather than myofiber dysfunction. We do note that single fiber force testing must be performed on permeabilized myofiber segments. While fresh, intact myofibers can be isolated from small murine muscles, like the lumbrical [79], the procedure is not technically feasible in larger muscles like the diaphragm (H. Westerblad, Karolinska Institutet, personal communication). As a consequence, the single myofiber approach tests the functional integrity of the sarcomere but may exclude assessment of the extrinsic regulation of excitation contraction coupling [80].

Despite these caveats, augmentation of intramuscular adipose depots likely plays a substantial role in diminution of diaphragm isometric force during overnutrition. For instance, in well-defined models of simultaneous intramuscular adiposity and contractile dysfunction such as chemical injury of the EDL, lipodystrophic mice unable to generate adipocytes are protected from post-injury isometric force deficits [37]. As we previously described in the obese diaphragm and again demonstrate here, the authors of the aforementioned study found that simple occupation of muscle cross-sectional area by adipocytes was insufficient to account for the measured isometric force reduction in wild type mice [37]. Together, these findings suggest that a negative impact of intramuscular adipose depots on contractile physiology may be exerted through disruption of normal tissue architecture or potentially via paracrine signaling to myofibers [81]. Clinical relevance is highlighted by reports linking reduced limb muscle density (computed tomography), indicative of elevated intramuscular adiposity, to reduced physical performance in elderly men [82], and even to impaired lung function in young adults with obesity [23].

In our study, *Thbs1*-dependent fibro-adipogenic remodeling and contractile dysfunction correspond with compromised diaphragm motion on non-invasive ultrasound—highlighting the manifestation of THBS1-driven changes in clinically measurable outcomes. Blockade of THBS1 has been shown to mitigate hyperglycemia-induced peritoneal fibrosis [83], while inhibition of THBS1-dependent TGFβ activation has been shown to reduce renal injury and proteinuria in mouse models of diabetic nephropathy [84]. To this end, pharmacological targeting of THBS1 and its downstream signaling pathways may also hold potential as a treatment for obesity-associated respiratory dysfunction.

## METHODS

### Animals

Wild type C57Bl/6J (Strain #000664) and *Thbs1^-/-^* mice (Strain #006141) were obtained from The Jackson Laboratory (Bar Harbor, ME). Jackson maintains *Thbs1^-/-^* mice on a C57Bl/6J background. Animals were housed in pathogen-free containment with a 12-hour light-dark cycle and *ad libitum* food and water. For DIO studies, mice received a normal chow diet (5L0D; LabDiet, St Louis, MO) until 2 months of age. CD-fed mice continued this for an additional 6 months, while HFD-fed mice switched to a diet containing 45% calories from lipid (D12451; Research Diets, New Brunswick, NJ) and subsequently maintained this for 6 months. Body composition was assessed via NMR (using the EchoMRI 4in1-500). Glucose tolerance testing was performed via intraperitoneal injection of a 10% dextrose solution (in sterile water) dosed at 1g/ kg total body weight. Glucose measurements were made using a One Touch glucometer (Lifespan, Milpitas, CA) on blood samples obtained from tail nicks at time points 15, 30, 60, 90 and 120 minutes after dextrose injection. The University of Michigan Institutional Animal Care and Use Committee (IACUC) approved all animal studies.

### Diaphragm Ultrasonography

Diaphragm ultrasonography was performed as previously described [68]. Briefly, diaphragms were localized by ultrasound (US) using a transversely oriented MS250 transducer (frequency 24 MHz) (Vevo 2100; Visual Sonics, Toronto, ON). Diaphragm motion, observed in M-mode, was recorded for 3 or more respiratory cycles. Excursion amplitude, inspiratory velocity, and expiratory velocity were measured on still images with values averaged over the recorded cycles.

### Ex vivo isometric force testing (muscle strips)

Isometric force testing on diaphragm strips was performed as previously described [27, 85]. Briefly, tetanic force was measured on 2- to 4-mm-wide muscle strips isolated from the mid-costal diaphragm. In a Krebs-Ringer bath containing 0.03 mmol/L tubocurarine chloride, held at 25°C and bubbled with 95% O2 and 5% CO2 (maintaining pH 7.4), an attached rib was sutured to a servomotor (model 305B; Aurora Scientific, Aurora, ON) and the free central tendon edge was sutured to a force transducer (model BG-50; Kulite Semiconductor Products, Leonia, NJ). The bath was electrically stimulated through a field generated between two platinum electrodes by a biphasic current stimulator (model 701A; Aurora Scientific). LabVIEW 2014 software (National Instruments, Austin, TX) controlled electrical pulse properties and servomotor activity while recording transducer data. Strips were adjusted to optimal length (Lo) at which a stimulus pulse elicited maximum isometric force (Po). Muscle cross-sectional area (CSA) was calculated using Lo and muscle mass. Specific force was calculated as the quotient of Po/CSA.

### Ex vivo isometric force testing (single myofibers)

Single myofiber contractility experiments were performed as previously described [86]. Costal diaphragm fiber bundles (4mm in length and 1mm in diameter) containing longitudinal arrays of myofibers were manually excised from the central region of the muscle then immediately immersed in ice-cold skinning solution [containing potassium propionate, imidazole and EGTA (Sigma)] for 30 minutes before storage at -80°C in a solution containing 50% glycerol by volume. Prior to each experiment, bundles were removed from storage and thawed before removal of individual fibers by manual extraction with fine forceps under a stereomicroscope.

Force responses and motor position were acquired through a 16-bit A-D board (NI-6052; National Instruments, Austin, TX) and analyzed on computer running custom-designed LabVIEW software (National Instruments). The solution-changing system (model 802A; Aurora Scientific) consisted of three glass-bottom chambers housed in a moveable, temperature-controlled stainless-steel plate. Movement of the plate relative to the fiber was achieved via two stepper motors: one to lower and raise the chamber array, and the other to translate the plate to a new chamber position.

Chamber 1 was filled with an EGTA-containing relaxing solution in which fibers could be manipulated. In this chamber, fibers were manually sutured to a servomotor (model 322; Aurora Scientific)-force transducer (model 403A; Aurora Scientific) apparatus with USP 10-0 monofilament nylon suture. Optimal sarcomere length was defined based on the diffraction pattern of laser light passed through the mounted fiber [87]. Once achieved, the corresponding optimal fiber length (Lf) was measured under a stereomicroscope. Fiber CSA was estimated (on fibers held at Lf) using width and depth measurements obtained from high-magnification digital images of top and side views of the fiber. Chambers 2 and 3 respectively contained low-[Ca2^+^] pre-activating solution and high-[Ca2^+^] activating solution. Fibers were exposed to Chamber 2 solution for a 3-minute priming period during which the passive force required to maintain the fiber at Lf was measured. Fibers were then transferred to Chamber 3 for elicitation of maximum isometric force (Fo) during sustained contraction. Maximum total isometric force was calculated as the difference between Fo and passive force. Specific force was calculated as the quotient of maximum total isometric force/ CSA.

### Single cell isolation

Diaphragmatic mononuclear cells were isolated through a protocol adapted from [28]. Costal diaphragms were excised, minced with scissors, and digested in collagenase type II (Worthington Biochemical, Lakewood, NJ) diluted to 0.067% by weight in serum-free DMEM (Thermo Fisher Scientific, Waltham, MA). After one 1-hour incubation at 37°C, samples were triturated 3-4 times through an 18-gauge needle, then incubated at 37°C for an additional 10 minutes. Next, collagenase was inactivated with an excess of DMEM containing 10% fetal bovine serum (FBS) by volume; and samples were sequentially passed through 100 μm and 40 μm cell strainers to remove debris. Erythrocyte lysis was achieved via 30-second exposure to hypotonic stress, after which cells were re-suspended in phosphate-buffered saline (PBS).

Flow cytometry analysis used established marker profiles [28]. Briefly, fresh cells were incubated in PBS with fluorophore-conjugated CD31, CD45, integrin α7, and Sca1 antibodies at dilutions indicated in (Supplementary Table 1) for 30 minutes at 4°C. DAPI was added for the final 5 minutes of the incubation to act as a dead cell marker. Cells were analyzed on a MoFlo Astrios EQ running Summit software (version 6.3; Beckman-Coulter, Brea, CA). Gates were established using fluorescence minus one approach; and plots were generated in FCS Express 7 (DeNovo Software). Cell isolation for scRNA-seq was identical; however, cells were stained only with DAPI, and all live mononuclear cells were isolated.

### Cell Culture

After isolation, FAPs were cultured for 4 days in 12-well plates containing standard medium: DMEM with 10% FBS and antibiotic/antimycotic [penicillin, streptomycin, and amphotericin B (Sigma)]. Cells were then seeded at 30% confluence in optical bottom 96-well plates with standard medium and allowed to attach over 24 hours. The medium was then changed to DMEM with 1% FBS +/-THBS1 5 μg/mL. After 3 days, cells were fixed, blocked, and immunostained as previously described [27] using antibody concentrations indicated in (Supplementary Table 1). For Ki67 staining, wash buffers and antibody diluents contained 0.2% Tween-20. Detergent was excluded for extracellular fibronectin staining. Counterstain with Alexa Fluor 488– conjugated phalloidin (Thermo Fisher Scientific) was used in specific experiments.

### Single cell RNA sequencing and bioinformatics analysis

Sequencing was performed by the University of Michigan Advanced Genomics Core, with libraries constructed and subsequently subjected to 151 paired end cycles on the NovaSeq-6000 platform (Illumina, San Diego, CA). Bcl2fastq2 Conversion Software (Illumina) was used to generate de-multiplexed Fastq files. Mapping and quantitation were also done by the Advanced Genomics Core, using the ENSEMBL GRCm38 reference, and Cell Ranger to generate feature-barcode matrices, and aggregate the results from different samples. Approximately 10 million reads were sequenced per sample.

To produce velocity plots, velocyto (v. 0.17.17) and scVelo (v. 0.0.4, with Python 3.7.12) were used. Velocyto was installed as a conda environment. For each sample, the t-SNE coordinates were exported from the Loupe Browser with their barcode. Then velocyto was run from the command line with the t-SNE coordinates for each sample, a gtf file containing positions of repetitive elements to mask, the position-sorted bam file of filtered raw reads, and the mouse gtf file “Mus musculus.GRCm38.98.gtf”. The output is a file in “Loom” format, designed to efficiently store single-cell datasets and metadata. The loom files were input to an R (v. 4.1.3) script with the package “reticulate” (v. 1.25), loaded to run Python in R. The Python package “scVelo” was imported to the R script to calculate the cellular dynamics. For each sample, using the t-SNE coordinates and the calculated velocity, with ggplot2 (v 2.3.4) library loaded, a t-SNE plot could be produced with the rate and direction streams. Using the Loupe Browser 5, differential expression was calculated within FAP subclusters, and comparing each sample versus the two others combined. This gave a table of p-values and fold-changes that were uploaded to iPathwayGuide (advaitabio.com), using a linear absolute fold change cutoff of 1.5. All three comparisons in a subcluster were combined as a meta-analysis in iPathwayGuide, to visualize Venn diagrams of the marker genes and corresponding pathways. Data were made publicly available through upload to NCI Gene Expression Omnibus (GSE241005). Accession numbers are as follows: GSM7713701 (WT CD), GSM7713702 (WT HFD), and GSM7713703 (KO HFD).

### Whole tissue gene expression profiling

Costal hemidiaphragms were cleaned of adherent tissues, snap-frozen in liquid nitrogen, and digested in Trizol (Thermo Fisher) with mechanical homogenization. Total RNA was isolated with RNEasy reagents (Qiagen, Germantown, MD) as per manufacturer’s protocol. QuantSeq 3’ mRNA sequencing (Lexogen, Vienna, Austria) was performed by the University of Michigan Advanced Genomics Core. Gene set enrichment analysis was performed using GSEA 4.1 software (University of California, San Diego). Volcano plots were generated in R Studio (Posit, PBC, Boston, MA). For quantitative PCR cDNA was synthesized with SuperScript II (Invitrogen, Carlsbad, CA), and the PCR reaction performed with SYBR Green (Thermo Fisher Scientific) on a StepOnePlus machine (Applied Biosystems). Primer sequences for indicated genes are as follows: *Lep* forward: CAGTGCCTATCCAGAAAGTC; reverse: ATCTTGGACAAACTCAGAATG. *Pdgfra* forward: TTGATGAAGGTGGAACTGCT; reverse: ATTCCTCTGCCTGACATTGAC. *Fn1* forward: CGTTCATCTCCACTTGAT; reverse: CAGTTGTGTGCTCCGATCTC. *Col3a1* forward: CTTCTGGTTCTCCTGGTC; reverse: CAACCTTCACCCTTATCTCC. *Emr1*forward: CTTTGGCTATGGGCTTCCAGTC; reverse: GCAAGGAGGACAGAGTTTATCGTG.

### Histological Analysis

Seven μm-thick formalin-fixed, paraffin-embedded sections of the costal hemidiaphragm were prepared as previously described [27] and included samples in both transverse and longitudinal planes with respect to myofiber orientation. Both sample types were approximately 2-4 mm wide and included tissue encompassing the entire rib to tendon extent of the costal diaphragm muscle. Longitudinal samples were analyzed along the entire rib-tendon length, while transverse sections were analyzed at the midpoint of the rib-tendon axis. Hematoxylin/ eosin (H/E) and picrosirius red staining were performed by standard methods. Fiber size measurements were made using transverse sections stained with fluorescein 405-conjugated wheat germ agglutinin (WGA) (Biotium) diluted 1:200 in HBSS and incubated for 30 minutes at room temperature. Immunohistochemistry for perilipin, THY1 and fibronectin was performed with primary-secondary antibody pairs as indicted in (Supplementary Table 1).

For myofiber typing analyses, excised costal diaphragm samples were sequentially submerged in 30% sucrose in PBS and the mixture of 30% sucrose in PBS/OCT (1:1) until the tissues no longer floated. Tissues were then placed in OCT solution and frozen in liquid nitrogen-cooled isopentane for cryosectioning. Seven μm-thick transverse cryosections were immunostained with two primary antibodies specifying type I and type IIa fibers (type IIb and type IIx fibers were unstained) as previously described [37]. Primary and secondary antibodies are indicated in (Supplementary Table 1).

For immunofluorescent staining of CD68 and PDGFRα, excised diaphragm tissues were directly embedded in OCT and quickly snap-frozen in liquid nitrogen-cooled isopentane. Seven μm-thick transverse or longitudinal cryosections (as described above) were fixed in 4% PFA/PBS for 5 min at room temperature, then blocked and permeabilized in 1% BSA/PBS or MOM blocking medium containing 0.5% Triton X-100. Tissue slides were stained with different combinations of primary antibodies in 1% BSA/PBS or MOM antibody medium containing 0.1% Triton X-100 overnight at 4’C, followed by corresponding secondary antibodies for 1 hour at room temperature. Primary-secondary antibody pairs are indicated in (Supplementary Table 1). Nuclei were counterstained with DAPI (diluted in deionized water) for 5 min at room temperature before mounting of samples Prolong Diamond (Invitrogen).

Samples were imaged using an Olympus DP72 camera mounted on an Olympus SZ61 microscope (Tokyo, Japan) or a Nikon A1 confocal microscope running NIS-Elements software (Olympus). For all forms of staining, at least 3 sections, separated from one another by at least 100 μm, were analyzed; and quantitative morphometry was performed using NIH ImageJ.

### Statistical Analysis

Statistical analysis was performed in GraphPad Prism 9 and employed Student’s t-test for two-group comparisons, one-way ANOVA (with Tukey post hoc test) for comparisons of two or more groups, two-way ANOVA (with Sidak post hoc test) for two variables, and linear regression for correlation analysis. P value <0.05 indicated statistical significance. Quantitative data are shown as mean +/- SD.

## AUTHOR CONTRIBUTIONS

E.D.B and T-H.C conceived of the study. E.D.B and T-H.C. designed the experiments with advice from D.R.C and S.V.B. E.D.B, M.S.W, R.K.V, S.H.K, C.S.D, and K.C.B performed the experiments. E.D.B and T-H.C analyzed the data with advice from S.V.B. E.D.B and T-H.C. wrote the manuscript and D.E.M and S.V.B edited the manuscript.

## ACKNOWLEDGEMENTS

We thank the University of Michigan Advanced Genomics Core for single cell and bulk RNA sequencing, the University of Michigan BCRF Bioinformatics Core for assistance with single cell analyses, and Dr. Jae-Eun Choi for assistance with bulk RNA sequencing analysis. We thank Dr. Helena Schotland for discussion on the clinical implications of our study. Research reported here was supported by the NHLBI (5 K08 HL14737701-05, to E.D.B.); NIDDK (R01DK095137 to T-H.C.); University of Michigan Bridging Fund, Biointerfaces Challenge Grant, and Michigan Integrative Musculoskeletal Health Core Center (MiMHC) P30 Center Pilot Feasibility Grant (to T-H.C); the Michigan Obesity Nutrition Research Center (AWD015548, to E.D.B.); and the Caswell Diabetes Institute of the University of Michigan (F052716, to E.D.B.). The Michigan Musculoskeletal Health Center Function Core, in which ultrasound experiments were performed, is supported by NIAMS under award number P30 AR069620. The University of Michigan Animal Phenotyping Core, in which body composition testing was performed, is supported by awards U2CDK135066, DK020572, and DK089503.

## REFERENCES

1. Hales, C.M., et al., Prevalence of Obesity and Severe Obesity Among Adults: United States, 2017-2018. NCHS Data Brief, 2020(360): p. 1-8.

2. Gibson, G.J., Obesity, respiratory function and breathlessness. Thorax, 2000. 55 Suppl 1(Suppl 1): p. S41–4.

3. Babb, T.G., et al., Dyspnea on exertion in obese women: association with an increased oxygen cost of breathing. Am J Respir Crit Care Med, 2008. 178(2): p. 116–23.

4. Whipp, B.J. and J.A. Davis, The ventilatory stress of exercise in obesity. Am Rev Respir Dis, 1984. 129(2 Pt 2): p. S90-2.

5. Sin, D.D., R.L. Jones, and S.F. Man, Obesity is a risk factor for dyspnea but not for airflow obstruction. Arch Intern Med, 2002. 162(13): p. 1477–81.

6. Kessler, R., et al., The obesity-hypoventilation syndrome revisited: a prospective study of 34 consecutive cases. Chest, 2001. 120(2): p. 369–76.

7. Mokhlesi, B., et al., Obesity hypoventilation syndrome: prevalence and predictors in patients with obstructive sleep apnea. Sleep Breath, 2007. 11(2): p. 117–24.

8. Macavei, V.M., et al., Diagnostic predictors of obesity-hypoventilation syndrome in patients suspected of having sleep disordered breathing. J Clin Sleep Med, 2013. 9(9): p. 879–84.

9. Laaban, J.P. and E. Chailleux, Daytime hypercapnia in adult patients with obstructive sleep apnea syndrome in France, before initiating nocturnal nasal continuous positive airway pressure therapy. Chest, 2005. 127(3): p. 710–5.

10. Banerjee, D., et al., Obesity hypoventilation syndrome: hypoxemia during continuous positive airway pressure. Chest, 2007. 131(6): p. 1678–84.

11. Lecube, A., et al., Asymptomatic sleep-disordered breathing in premenopausal women awaiting bariatric surgery. Obes Surg, 2010. 20(4): p. 454–61.

12. Bernhardt, V., et al., Aerobic exercise training without weight loss reduces dyspnea on exertion in obese women. Respir Physiol Neurobiol, 2016. 221: p. 64–70.

13. Bernhardt, V., et al., Weight loss reduces dyspnea on exertion and unpleasantness of dyspnea in obese men. Respir Physiol Neurobiol, 2019. 261: p. 55–61.

14. Marines-Price, R., et al., Dyspnea on exertion provokes unpleasantness and negative emotions in women with obesity. Respir Physiol Neurobiol, 2019. 260: p. 131–136.

15. Castro-Anon, O., et al., Obesity-hypoventilation syndrome: increased risk of death over sleep apnea syndrome. PLoS One, 2015. 10(2): p. e0117808.

16. De Jong, A., G. Chanques, and S. Jaber, Mechanical ventilation in obese ICU patients: from intubation to extubation. Crit Care, 2017. 21(1): p. 63.

17. Scano, G., L. Stendardi, and G.I. Bruni, The respiratory muscles in eucapnic obesity: their role in dyspnea. Respir Med, 2009. 103(9): p. 1276–85.

18. Cherniack, R.M. and C.A. Guenter, The efficiency of the respiratory muscles in obesity. Can J Biochem Physiol, 1961. 39: p. 1215–22.

19. Sahebjami, H., Dyspnea in obese healthy men. Chest, 1998. 114(5): p. 1373–7.

20. Chlif, M., et al., Inspiratory muscle activity during incremental exercise in obese men. Int J Obes (Lond), 2007. 31(9): p. 1456–63.

21. Lennmarken, C., et al., Skeletal muscle function and metabolism in obese women. JPEN J Parenter Enteral Nutr, 1986. 10(6): p. 583–7.

22. Newham, D.J., et al., The strength, contractile properties and radiological density of skeletal muscle before and 1 year after gastroplasty. Clin Sci (Lond), 1988. 74(1): p. 79–83.

23. Yu, X., et al., Association of Muscle Fat Content and Muscle Mass With Impaired Lung Function in Young Adults With Obesity: Evaluation With MRI. Acad Radiol, 2023.

24. Fadell, E.J., et al., Fatty infiltration of respiratory muscles in the Pick-wickian syndrome. N Engl J Med, 1962. 266: p. 861–3.

25. Popkin, B.M., et al., Individuals with obesity and COVID-19: A global perspective on the epidemiology and biological relationships. Obes Rev, 2020. 21(11): p. e13128.

26. Shi, Z., et al., Diaphragm Pathology in Critically Ill Patients With COVID-19 and Postmortem Findings From 3 Medical Centers. JAMA Intern Med, 2021. 181(1): p. 122–124.

27. Buras, E.D., et al., Fibro-Adipogenic Remodeling of the Diaphragm in Obesity-Associated Respiratory Dysfunction. Diabetes, 2019. 68(1): p. 45–56.

28. Joe, A.W., et al., Muscle injury activates resident fibro/adipogenic progenitors that facilitate myogenesis. Nat Cell Biol, 2010. 12(2): p. 153–63.

29. Uezumi, A., et al., Mesenchymal progenitors distinct from satellite cells contribute to ectopic fat cell formation in skeletal muscle. Nat Cell Biol, 2010. 12(2): p. 143–52.

30. Uezumi, A., et al., Identification and characterization of PDGFRalpha+ mesenchymal progenitors in human skeletal muscle. Cell Death Dis, 2014. 5(4): p. e1186.

31. Heredia, J.E., et al., Type 2 innate signals stimulate fibro/adipogenic progenitors to facilitate muscle regeneration. Cell, 2013. 153(2): p. 376–88.

32. Lemos, D.R., et al., Nilotinib reduces muscle fibrosis in chronic muscle injury by promoting TNF-mediated apoptosis of fibro/adipogenic progenitors. Nat Med, 2015. 21(7): p. 786–94.

33. Wosczyna, M.N., et al., Mesenchymal Stromal Cells Are Required for Regeneration and Homeostatic Maintenance of Skeletal Muscle. Cell Rep, 2019. 27(7): p. 2029–2035 e5.

34. Uezumi, A., et al., Fibrosis and adipogenesis originate from a common mesenchymal progenitor in skeletal muscle. J Cell Sci, 2011. 124(Pt 21): p. 3654–64.

35. Mozzetta, C., et al., Fibroadipogenic progenitors mediate the ability of HDAC inhibitors to promote regeneration in dystrophic muscles of young, but not old Mdx mice. EMBO Mol Med, 2013. 5(4): p. 626–39.

36. Larouche, J.A., et al., Spatiotemporal mapping of immune and stem cell dysregulation after volumetric muscle loss. JCI Insight, 2023.

37. Biltz, N.K., et al., Infiltration of intramuscular adipose tissue impairs skeletal muscle contraction. J Physiol, 2020. 598(13): p. 2669–2683.

38. McKellar, D.W., et al., Large-scale integration of single-cell transcriptomic data captures transitional progenitor states in mouse skeletal muscle regeneration. Commun Biol, 2021. 4(1): p. 1280.

39. Farup, J., et al., Human skeletal muscle CD90(+) fibro-adipogenic progenitors are associated with muscle degeneration in type 2 diabetic patients. Cell Metab, 2021. 33(11): p. 2201–2214 e11.

40. Varma, V., et al., Thrombospondin-1 is an adipokine associated with obesity, adipose inflammation, and insulin resistance. Diabetes, 2008. 57(2): p. 432–9.

41. Ramis, J.M., et al., Carboxypeptidase E and thrombospondin-1 are differently expressed in subcutaneous and visceral fat of obese subjects. Cell Mol Life Sci, 2002. 59(11): p. 1960–71.

42. Kong, P., et al., Thrombospondin-1 regulates adiposity and metabolic dysfunction in diet-induced obesity enhancing adipose inflammation and stimulating adipocyte proliferation. Am J Physiol Endocrinol Metab, 2013. 305(3): p. E439–50.

43. Matsuo, Y., et al., Thrombospondin 1 as a novel biological marker of obesity and metabolic syndrome. Metabolism, 2015. 64(11): p. 1490–9.

44. Inoue, M., et al., Thrombospondin 1 mediates high-fat diet-induced muscle fibrosis and insulin resistance in male mice. Endocrinology, 2013. 154(12): p. 4548–59.

45. Belotti, D., et al., Thrombospondin-1 promotes mesenchymal stromal cell functions via TGFbeta and in cooperation with PDGF. Matrix Biol, 2016. 55: p. 106–116.

46. De Luna, N., et al., Role of thrombospondin 1 in macrophage inflammation in dysferlin myopathy. J Neuropathol Exp Neurol, 2010. 69(6): p. 643–53.

47. Suarez-Calvet, X., et al., Thrombospondin-1 mediates muscle damage in brachio-cervical inflammatory myopathy and systemic sclerosis. Neurol Neuroimmunol Neuroinflamm, 2020. 7(3).

48. Madaro, L., et al., Denervation-activated STAT3-IL-6 signalling in fibro-adipogenic progenitors promotes myofibres atrophy and fibrosis. Nat Cell Biol, 2018. 20(8): p. 917–927.

49. Malecova, B., et al., Dynamics of cellular states of fibro-adipogenic progenitors during myogenesis and muscular dystrophy. Nat Commun, 2018. 9(1): p. 3670.

50. Giuliani, G., M. Rosina, and A. Reggio, Signaling pathways regulating the fate of fibro/adipogenic progenitors (FAPs) in skeletal muscle regeneration and disease. FEBS J, 2022. 289(21): p. 6484–6517.

51. Rubenstein, A.B., et al., Single-cell transcriptional profiles in human skeletal muscle. Sci Rep, 2020. 10(1): p. 229.

52. Hepler, C., et al., Identification of functionally distinct fibro-inflammatory and adipogenic stromal subpopulations in visceral adipose tissue of adult mice. Elife, 2018. 7.

53. Fitzgerald, G., et al., MME(+) fibro-adipogenic progenitors are the dominant adipogenic population during fatty infiltration in human skeletal muscle. Commun Biol, 2023. 6(1): p. 111.

54. Merrick, D., et al., Identification of a mesenchymal progenitor cell hierarchy in adipose tissue. Science, 2019. 364(6438).

55. Petrus, P., et al., Specific loss of adipocyte CD248 improves metabolic health via reduced white adipose tissue hypoxia, fibrosis and inflammation. EBioMedicine, 2019. 44: p. 489–501.

56. Egea, V., et al., Tissue inhibitor of metalloproteinase-1 (TIMP-1) regulates mesenchymal stem cells through let-7f microRNA and Wnt/beta-catenin signaling. Proc Natl Acad Sci U S A, 2012. 109(6): p. E309–16.

57. Lee, E.J., et al., Fibromodulin: a master regulator of myostatin controlling progression of satellite cells through a myogenic program. FASEB J, 2016. 30(8): p. 2708–19.

58. Descamps, S., et al., Inhibition of myoblast differentiation by Sfrp1 and Sfrp2. Cell Tissue Res, 2008. 332(2): p. 299–306.

59. Giordani, L., et al., High-Dimensional Single-Cell Cartography Reveals Novel Skeletal Muscle-Resident Cell Populations. Mol Cell, 2019. 74(3): p. 609–621 e6.

60. Docheva, D., et al., Tenomodulin is necessary for tenocyte proliferation and tendon maturation. Mol Cell Biol, 2005. 25(2): p. 699–705.

61. Yao, L., et al., Paracrine signalling during ZEB1-mediated epithelial-mesenchymal transition augments local myofibroblast differentiation in lung fibrosis. Cell Death Differ, 2019. 26(5): p. 943–957.

62. Meng, Q., et al., Myofibroblast-Specific TGFbeta Receptor II Signaling in the Fibrotic Response to Cardiac Myosin Binding Protein C-Induced Cardiomyopathy. Circ Res, 2018. 123(12): p. 1285–1297.

63. Qadri, M., et al., Proteoglycan-4 regulates fibroblast to myofibroblast transition and expression of fibrotic genes in the synovium. Arthritis Res Ther, 2020. 22(1): p. 113.

64. Lerbs, T., et al., CD47 prevents the elimination of diseased fibroblasts in scleroderma. JCI Insight, 2020. 5(16).

65. Dawson, D.W., et al., CD36 mediates the In vitro inhibitory effects of thrombospondin-1 on endothelial cells. J Cell Biol, 1997. 138(3): p. 707–17.

66. Yadav, H., et al., Protection from obesity and diabetes by blockade of TGF-beta/Smad3 signaling. Cell Metab, 2011. 14(1): p. 67–79.

67. Jaitin, D.A., et al., Lipid-Associated Macrophages Control Metabolic Homeostasis in a Trem2-Dependent Manner. Cell, 2019. 178(3): p. 686–698 e14.

68. Whitehead, N.P., et al., Validation of ultrasonography for non-invasive assessment of diaphragm function in muscular dystrophy. J Physiol, 2016. 594(24): p. 7215–7227.

69. McLaren, K.M., Immunohistochemical localisation of thrombospondin in human megakaryocytes and platelets. J Clin Pathol, 1983. 36(2): p. 197–9.

70. Jaffe, E.A., et al., Cultured human fibroblasts synthesize and secrete thrombospondin and incorporate it into extracellular matrix. Proc Natl Acad Sci U S A, 1983. 80(4): p. 998–1002.

71. Mosher, D.F., M.J. Doyle, and E.A. Jaffe, Synthesis and secretion of thrombospondin by cultured human endothelial cells. J Cell Biol, 1982. 93(2): p. 343–8.

72. Isenberg, J.S. and D.D. Roberts, Thrombospondin-1 in maladaptive aging responses: a concept whose time has come. Am J Physiol Cell Physiol, 2020. 319(1): p. C45–C63.

73. von Toerne, C., et al., MASP1, THBS1, GPLD1 and ApoA-IV are novel biomarkers associated with prediabetes: the KORA F4 study. Diabetologia, 2016. 59(9): p. 1882-92.

74. Bohm, A., et al., TGF-beta Contributes to Impaired Exercise Response by Suppression of Mitochondrial Key Regulators in Skeletal Muscle. Diabetes, 2016. 65(10): p. 2849–61.

75. Saito, Y., et al., Exercise enhances skeletal muscle regeneration by promoting senescence in fibro-adipogenic progenitors. Nat Commun, 2020. 11(1): p. 889.

76. Murphy-Ullrich, J.E., Thrombospondin 1 and Its Diverse Roles as a Regulator of Extracellular Matrix in Fibrotic Disease. J Histochem Cytochem, 2019. 67(9): p. 683–699.

77. Hocevar, B.A., T.L. Brown, and P.H. Howe, TGF-beta induces fibronectin synthesis through a c-Jun N-terminal kinase-dependent, Smad4-independent pathway. EMBO J, 1999. 18(5): p. 1345-56.

78. Salimena, M.C., J. Lagrota-Candido, and T. Quirico-Santos, Gender dimorphism influences extracellular matrix expression and regeneration of muscular tissue in mdx dystrophic mice. Histochem Cell Biol, 2004. 122(5): p. 435–44.

79. Claflin, D.R. and S.V. Brooks, Direct observation of failing fibers in muscles of dystrophic mice provides mechanistic insight into muscular dystrophy. Am J Physiol Cell Physiol, 2008. 294(2): p. C651–8.

80. Roche, S.M., et al., Measurement of Maximum Isometric Force Generated by Permeabilized Skeletal Muscle Fibers. J Vis Exp, 2015(100): p. e52695.

81. Pellegrinelli, V., et al., Human Adipocytes Induce Inflammation and Atrophy in Muscle Cells During Obesity. Diabetes, 2015. 64(9): p. 3121–34.

82. Orwoll, E.S., et al., CT Muscle Density, D3Cr Muscle Mass, and Body Fat Associations With Physical Performance, Mobility Outcomes, and Mortality Risk in Older Men. J Gerontol A Biol Sci Med Sci, 2022. 77(4): p. 790–799.

83. Jiang, N., et al., Blockade of thrombospondin-1 ameliorates high glucose-induced peritoneal fibrosis through downregulation of TGF-beta1/Smad3 signaling pathway. J Cell Physiol, 2020. 235(1): p. 364–379.

84. Lu, A., et al., Blockade of TSP1-dependent TGF-beta activity reduces renal injury and proteinuria in a murine model of diabetic nephropathy. Am J Pathol, 2011. 178(6): p. 2573–86.

85. Brooks, S.V. and J.A. Faulkner, Contractile properties of skeletal muscles from young, adult and aged mice. J Physiol, 1988. 404: p. 71–82.

86. Larkin, L.M., et al., Weakness of whole muscles in mice deficient in Cu, Zn superoxide dismutase is not explained by defects at the level of the contractile apparatus. Age (Dordr), 2013. 35(4): p. 1173–81.

87. Mendias, C.L., et al., Decreased specific force and power production of muscle fibers from myostatin-deficient mice are associated with a suppression of protein degradation. J Appl Physiol (1985), 2011. 111(1): p. 185-91.

