## Supplementary Information for "Thrombospondin-1 promotes fibro-adipogenic stromal expansion and contractile dysfunction of the diaphragm in obesity"

### Figure S1: Diaphragm mononuclear cell annotation, FAP1 marker genes

A

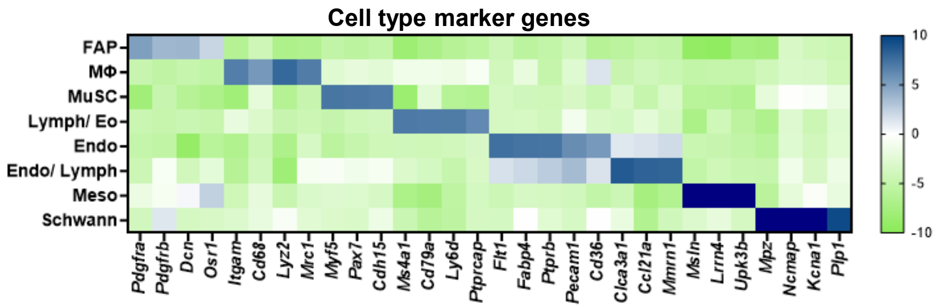


B

| Gene | Average | Log2 FC | p-val |
|---|---|---|---|
| *Cxcl14* | 12.85 | 4.01 | 2.17E-43 |
| *Hsd11b1* | 2.97 | 2.61 | 3.17E-18 |
| *Vtn* | 1.73 | 2.69 | 3.25E-18 |
| *Kcnk2* | 1.06 | 2.45 | 3.73E-15 |
| *Apod* | 6.31 | 2.72 | 1.03E-14 |
| *Crispld2* | 3.38 | 2.37 | 1.23E-14 |
| *Plau* | 1.92 | 2.37 | 1.45E-14 |
| *Ccl11* | 6.20 | 2.39 | 1.62E-14 |
| *Ces1d* | 1.80 | 2.31 | 9.48E-14 |
| *Gask1b* | 1.33 | 2.24 | 8.58E-13 |
| *G0s2* | 2.66 | 2.19 | 6.05E-12 |
| *Lifr* | 1.61 | 2.14 | 1.09E-11 |
| *Adm* | 2.89 | 2.08 | 8.72E-11 |
| *Smoc2* | 14.41 | 1.97 | 7.16E-10 |
| *Inmt* | 4.84 | 1.95 | 1.28E-09 |
| *Vwa1* | 1.62 | 1.93 | 2.62E-09 |
| *Col15a1* | 7.66 | 1.89 | 4.33E-09 |
| *Abca6* | 1.09 | 1.92 | 4.41E-09 |
| *Gpm6b* | 1.30 | 1.86 | 1.69E-08 |
| *Itih5* | 6.44 | 1.82 | 2.22E-08 |
| *Col6a6* | 1.64 | 1.83 | 2.60E-08 |
| *Lamb1* | 2.95 | 1.77 | 7.40E-08 |
| *Rgmb* | 1.03 | 1.78 | 7.53E-08 |
| *Col6a3* | 5.23 | 1.75 | 9.89E-08 |
| *Scn7a* | 2.07 | 1.71 | 2.89E-07 |
| *Atp1a2* | 1.08 | 1.69 | 5.81E-07 |
| *Abca8b* | 2.98 | 1.64 | 1.02E-06 |
| *Abca8a* | 9.88 | 1.63 | 1.07E-06 |
| *Phlda1* | 9.31 | 1.62 | 1.40E-06 |
| *Iigp1* | 1.62 | 1.66 | 1.65E-06 |
| *Cxcl1* | 8.23 | 1.59 | 4.16E-06 |
| *Spry1* | 1.32 | 1.58 | 4.33E-06 |
| *Podn* | 1.84 | 1.56 | 4.95E-06 |
| *Ms4a4d* | 1.80 | 1.55 | 5.68E-06 |
| *Thbs1* | 3.13 | 1.60 | 5.99E-06 |
| *Dpep1* | 4.40 | 1.54 | 7.36E-06 |
| *Col5a3* | 2.45 | 1.54 | 8.19E-06 |
| *Bmp4* | 1.77 | 1.54 | 9.00E-06 |
| *Sparcl1* | 6.63 | 1.52 | 1.08E-05 |
| *Sdc2* | 4.39 | 1.51 | 1.15E-05 |
| *Lum* | 16.06 | 1.50 | 1.38E-05 |
| *Hspg2* | 3.99 | 1.50 | 1.43E-05 |
| *Oaf* | 3.47 | 1.47 | 2.38E-05 |
| *Col4a1* | 6.54 | 1.45 | 3.01E-05 |
| *Lama2* | 4.26 | 1.45 | 3.13E-05 |
| *Meox1* | 1.21 | 1.47 | 3.71E-05 |
| *F3* | 1.24 | 1.46 | 4.06E-05 |
| *Serpine1* | 1.12 | 1.46 | 4.77E-05 |
| *Nnmt* | 1.01 | 1.41 | 7.28E-05 |
| *Fibin* | 4.34 | 1.40 | 9.18E-05 |

C

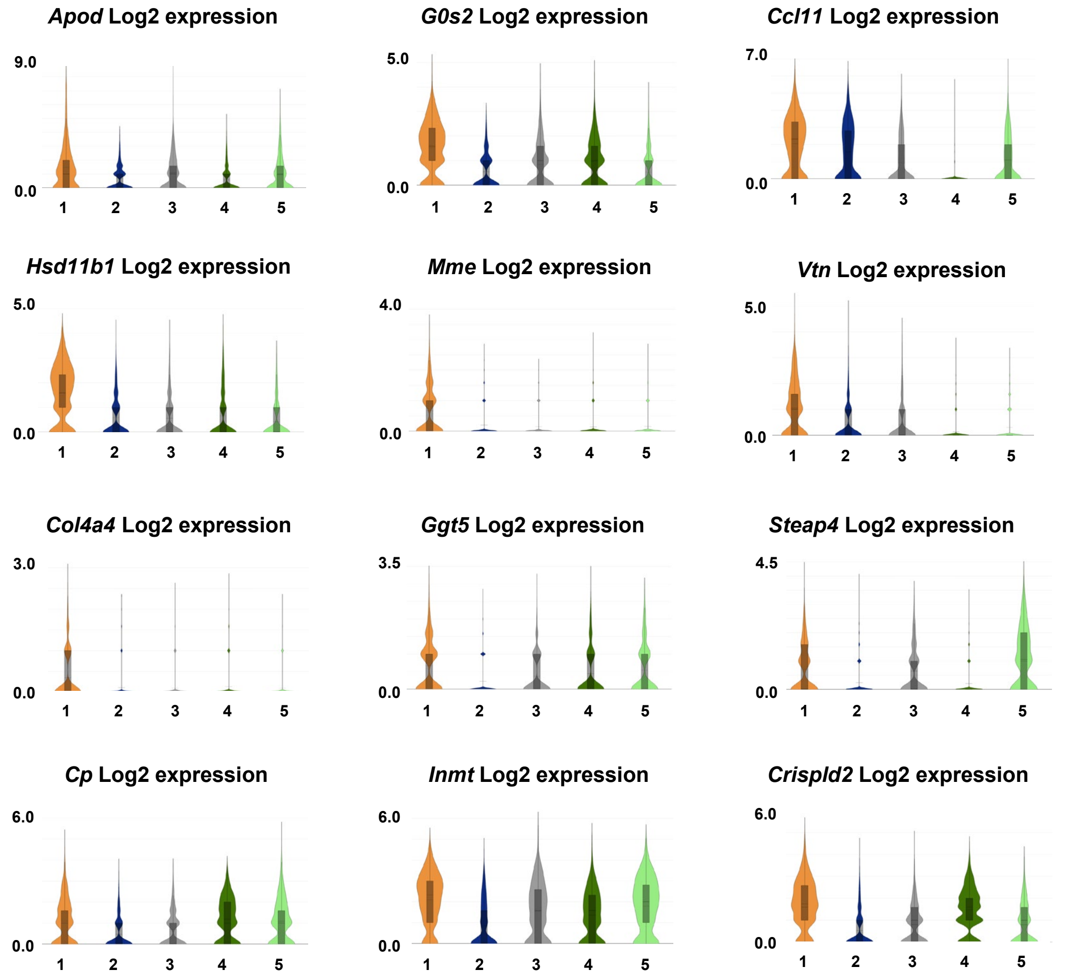


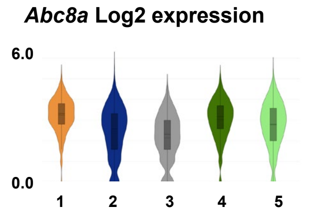


D

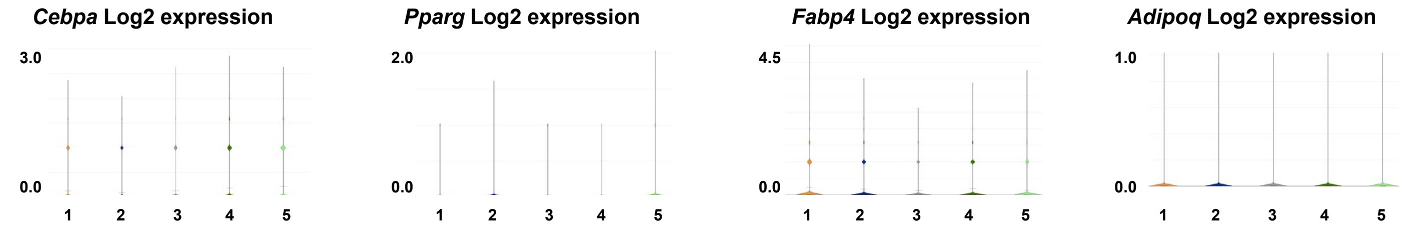


**Figure S1. (A)** Heat map demonstrating cell-specific gene expression used for diaphragm mononuclear cell annotation. **(B)** FAP1 sub-population: Top 50 enriched genes. **(C)** Violin plots demonstrating expression of adipocyte precursor genes in FAP sub-populations. **(D)** Violin plots demonstrating limited expression of committed preadipocyte and adipocyte markers in FAP sub-populations. FAP1, FAP2, FAP3, FAP4 and FAP5 respectively labeled 1, 2, 3, 4, 5.

### Figure S2: FAP2 marker genes

**A**

| Gene | Average | Log2 FC | p-val |
|---|---|---|---|
| *Stmn4* | 1.25 | 3.62 | 1.36E-40 |
| *Efhd1* | 1.09 | 3.60 | 1.36E-40 |
| *Sema3c* | 2.70 | 3.17 | 1.23E-31 |
| *Ccn3* | 1.45 | 3.05 | 1.78E-28 |
| *Sbsn* | 1.11 | 2.85 | 7.31E-22 |
| *Cd55* | 6.85 | 2.75 | 2.87E-24 |
| *Fbn1* | 16.62 | 2.70 | 1.85E-23 |
| *Pla1a* | 2.51 | 2.67 | 1.99E-22 |
| *Thy1* | 1.06 | 2.39 | 3.26E-17 |
| *Uap1* | 4.24 | 2.38 | 8.13E-18 |
| *Cadm3* | 1.04 | 2.35 | 6.58E-17 |
| *Dbn1* | 1.74 | 2.31 | 1.50E-16 |
| *Cd248* | 6.26 | 2.28 | 2.46E-16 |
| *Gfpt2* | 4.05 | 2.22 | 2.59E-15 |
| *Adgrd1* | 1.54 | 2.22 | 3.68E-15 |
| *Ugdh* | 2.95 | 2.20 | 7.35E-15 |
| *Dpp4* | 1.39 | 2.18 | 1.84E-14 |
| *Pcolce2* | 5.08 | 2.11 | 8.04E-14 |
| *Vcan* | 3.01 | 2.09 | 1.61E-13 |
| *Anxa3* | 3.88 | 2.09 | 1.60E-13 |
| *Mustn1* | 1.34 | 2.08 | 6.58E-13 |
| *Mfap5* | 26.85 | 2.08 | 1.68E-13 |
| *Zfp385a* | 1.27 | 2.01 | 2.50E-12 |
| *Pi16* | 29.97 | 1.97 | 4.39E-12 |
| *Fstl1* | 15.37 | 1.96 | 5.11E-12 |
| *Tmem100* | 3.08 | 1.96 | 9.55E-12 |
| *Pcsk6* | 6.32 | 1.96 | 7.44E-12 |
| *Creb5* | 2.08 | 1.94 | 1.51E-11 |
| *Adamts5* | 4.98 | 1.90 | 4.16E-11 |
| *Has1* | 1.89 | 1.87 | 5.86E-10 |
| *Ugp2* | 3.38 | 1.84 | 2.02E-10 |
| *Axl* | 4.52 | 1.81 | 4.48E-10 |
| *Ackr3* | 5.79 | 1.80 | 6.52E-10 |
| *Aspn* | 11.72 | 1.79 | 9.45E-10 |
| *Emilin2* | 2.27 | 1.74 | 3.71E-09 |
| *Fndc1* | 4.34 | 1.72 | 4.92E-09 |
| *Efemp1* | 3.33 | 1.71 | 8.77E-09 |
| *Postn* | 1.78 | 1.66 | 7.10E-07 |
| *Prss23* | 1.81 | 1.66 | 3.60E-08 |
| *Hspb8* | 1.11 | 1.62 | 8.55E-08 |
| *Thbd* | 1.39 | 1.60 | 1.25E-07 |
| *Loxl2* | 1.61 | 1.58 | 1.65E-07 |
| *Lrrn4cl* | 1.51 | 1.58 | 1.77E-07 |
| *Cd34* | 9.65 | 1.57 | 1.40E-07 |
| *Ddr2* | 2.22 | 1.55 | 3.27E-07 |
| *Tnxb* | 9.14 | 1.52 | 4.57E-07 |
| *Bmp1* | 1.68 | 1.52 | 6.01E-07 |
| *Ahnak2* | 1.81 | 1.52 | 6.32E-07 |
| *Rhoq* | 1.61 | 1.51 | 7.25E-07 |
| *Tppp3* | 7.58 | 1.49 | 9.38E-07 |

**B**

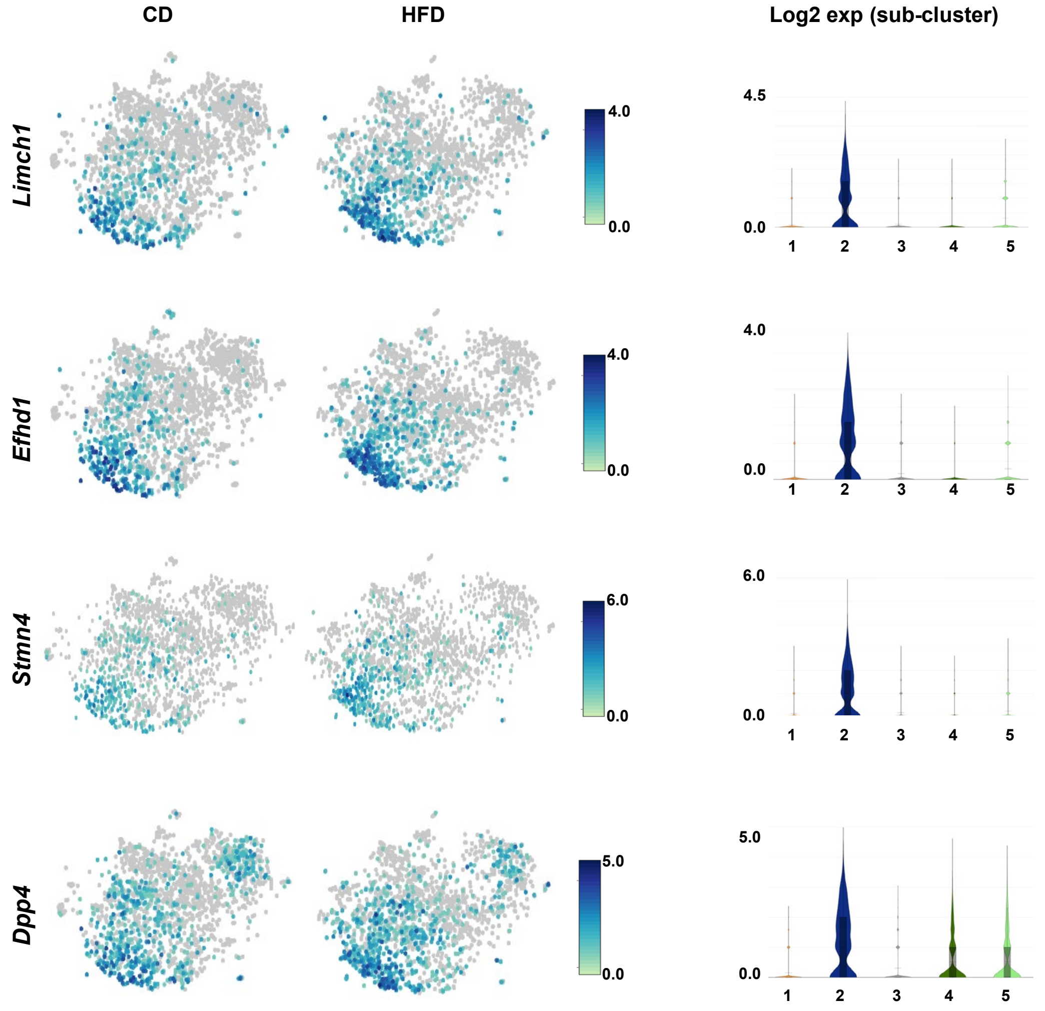


**C**

**D**

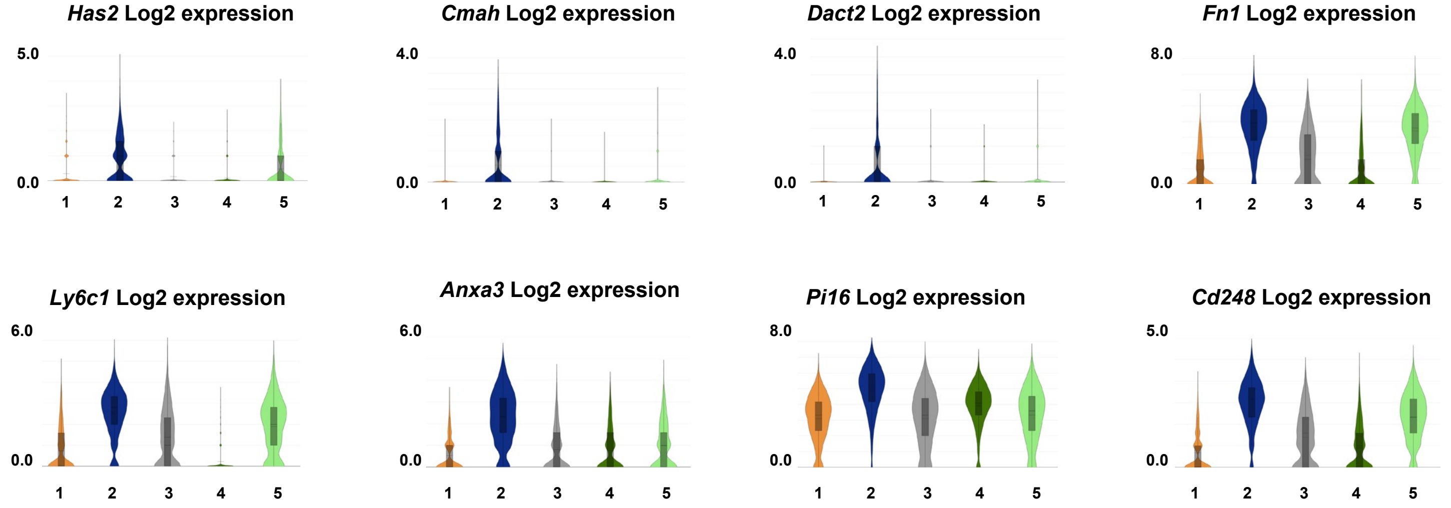


**Figure S2. (A)** FAP2 sub-population: Top 50 enriched genes. **(B)** FAP2-enriched genes with tSNE plots demonstrating expression in 6-month control diet (CD) versus high fat diet (HFD)-fed FAPs. Color gradient indicates Log2 expression. Violin plots show FAP sub-population-specific expression of the same genes. **(C)** Violin plots indicating sub-population-specific expression of genes previously associated with adipose fibroinflammatory progenitors (FIPs). **(D)** Violin plots indicating sub-population-specific expression of fibronectin (*Fn1*) and endosialin (CD248) (*Cd248*). FAP1, FAP2, FAP3, FAP4 and FAP5 respectively labeled 1, 2, 3, 4, 5.

### Figure S3: Marker genes: FAP3, FAP4, FAP5 subclusters

**A**

| Gene | Average | Log2 FC | p-val |
|---|---|---|---|
| *Timp1* | 1.10 | 1.77 | 1.79E-05 |
| *Sult1e1* | 0.94 | 1.58 | 2.68E-04 |
| *Col6a5* | 2.15 | 1.52 | 1.10E-04 |
| *Sfrp4* | 1.78 | 1.52 | 3.60E-04 |
| *Penk* | 2.50 | 1.49 | 8.34E-05 |
| *Apod* | 3.70 | 1.48 | 1.34E-03 |
| *Spon2* | 0.91 | 1.37 | 7.81E-04 |
| *Rbp1* | 5.95 | 1.31 | 9.33E-04 |
| *Igfbp7* | 17.77 | 1.30 | 1.01E-03 |
| *Serpinf1* | 21.00 | 1.28 | 1.22E-03 |
| *Sfrp1* | 2.24 | 1.28 | 1.63E-03 |
| *C7* | 2.55 | 1.27 | 1.87E-03 |
| *Phlda3* | 1.74 | 1.25 | 1.93E-03 |
| *Pcolce* | 8.22 | 1.23 | 2.23E-03 |
| *Cygb* | 4.85 | 1.22 | 2.68E-03 |
| *Steap4* | 0.89 | 1.22 | 3.76E-03 |
| *Mgp* | 24.54 | 1.21 | 3.54E-03 |
| *Cst3* | 44.70 | 1.21 | 3.50E-03 |
| *Olfml3* | 2.03 | 1.21 | 3.38E-03 |
| *Mmp23* | 2.31 | 1.18 | 4.52E-03 |
| *Il11ra1* | 5.13 | 1.18 | 4.23E-03 |
| *Mfap2* | 0.82 | 1.17 | 5.88E-03 |
| *Gpx3* | 10.32 | 1.17 | 4.98E-03 |
| *Inmt* | 3.30 | 1.17 | 5.62E-03 |
| *Igfbp4* | 13.73 | 1.15 | 5.95E-03 |
| *Gstm2* | 1.65 | 1.14 | 7.05E-03 |
| *Nbl1* | 7.90 | 1.13 | 7.62E-03 |
| *Mustn1* | 0.91 | 1.11 | 1.25E-02 |
| *Gpx8* | 1.65 | 1.11 | 1.00E-02 |
| *Selenom* | 7.40 | 1.11 | 9.43E-03 |
| *Sod3* | 1.79 | 1.11 | 1.07E-02 |
| *Entpd2* | 4.39 | 1.10 | 1.05E-02 |
| *Fbln1* | 4.68 | 1.09 | 1.27E-02 |
| *F3* | 1.03 | 1.07 | 1.74E-02 |
| *Lbp* | 2.14 | 1.07 | 1.60E-02 |
| *Lysmd2* | 1.22 | 1.06 | 1.67E-02 |
| *Gsn* | 147.09 | 1.04 | 1.89E-02 |
| *Mgst1* | 3.38 | 1.04 | 1.94E-02 |
| *Nenf* | 4.91 | 1.04 | 2.00E-02 |
| *Pdlim2* | 1.97 | 1.04 | 2.16E-02 |
| *Csrp2* | 3.33 | 1.03 | 2.32E-02 |
| *Ctsk* | 1.38 | 1.02 | 2.53E-02 |
| *Dpep1* | 3.37 | 1.02 | 2.56E-02 |
| *Tmed3* | 3.81 | 1.01 | 2.66E-02 |
| *Tmem100* | 2.07 | 1.01 | 3.00E-02 |
| *Vkorc1* | 2.84 | 1.01 | 2.74E-02 |
| *Serping1* | 16.48 | 1.00 | 2.84E-02 |
| *Maged2* | 1.81 | 1.00 | 3.21E-02 |
| *Bgn* | 14.18 | 0.99 | 3.04E-02 |
| *Cd302* | 4.76 | 0.98 | 3.47E-02 |

**B**

| Gene | Average | Log2 FC | p-val |
|---|---|---|---|
| *Col6a5* | 4.05 | 2.96 | 2.81E-20 |
| *Sult1e1* | 1.48 | 2.55 | 2.08E-12 |
| *Cyp1b1* | 1.04 | 2.45 | 5.67E-14 |
| *Has1* | 2.37 | 2.16 | 2.27E-10 |
| *Cxcl13* | 1.66 | 2.10 | 3.15E-04 |
| *Steap4* | 1.31 | 1.99 | 5.99E-09 |
| *C3* | 28.98 | 1.98 | 3.40E-09 |
| *Fst* | 2.18 | 1.87 | 8.72E-08 |
| *Ptgs2* | 2.79 | 1.80 | 4.87E-07 |
| *Ifi204* | 2.10 | 1.76 | 4.51E-07 |
| *C4b* | 2.60 | 1.76 | 2.09E-06 |
| *Tnfaip6* | 3.80 | 1.68 | 2.16E-06 |
| *Scara3* | 1.10 | 1.67 | 2.87E-06 |
| *Myc* | 1.85 | 1.53 | 3.17E-05 |
| *Mmp14* | 1.29 | 1.53 | 3.19E-05 |
| *C1s1* | 5.99 | 1.53 | 2.35E-05 |
| *Lgi2* | 1.38 | 1.51 | 4.22E-05 |
| *Ifi205* | 2.86 | 1.47 | 8.16E-05 |
| *Aebp1* | 9.97 | 1.46 | 7.35E-05 |
| *Steap3* | 1.84 | 1.45 | 1.06E-04 |
| *Ebf2* | 1.83 | 1.44 | 1.27E-04 |
| *Ddr2* | 2.24 | 1.43 | 1.33E-04 |
| *Penk* | 2.35 | 1.38 | 3.56E-04 |
| *Slit3* | 1.46 | 1.37 | 3.60E-04 |
| *Cdon* | 1.37 | 1.35 | 4.48E-04 |
| *Pdgfra* | 4.09 | 1.32 | 5.74E-04 |
| *Ugdh* | 2.13 | 1.32 | 7.44E-04 |
| *Hk2* | 1.87 | 1.32 | 8.52E-04 |
| *Scara5* | 3.13 | 1.32 | 6.51E-04 |
| *Man2a1* | 2.34 | 1.30 | 7.94E-04 |
| *Gfpt2* | 2.88 | 1.30 | 8.68E-04 |
| *Ifi211* | 1.64 | 1.30 | 8.45E-04 |
| *Cd248* | 4.30 | 1.28 | 1.13E-03 |
| *C1ra* | 3.56 | 1.27 | 1.23E-03 |
| *Igf1* | 4.43 | 1.27 | 1.34E-03 |
| *Lox* | 1.60 | 1.26 | 1.80E-03 |
| *Tshz2* | 2.21 | 1.21 | 2.55E-03 |
| *Arhgap20* | 1.04 | 1.21 | 2.94E-03 |
| *Lrp1* | 10.37 | 1.20 | 2.85E-03 |
| *Gpc3* | 2.30 | 1.19 | 3.84E-03 |
| *Pdgfrb* | 1.06 | 1.18 | 4.08E-03 |
| *Ace* | 2.25 | 1.18 | 3.99E-03 |
| *Ar* | 1.90 | 1.17 | 4.10E-03 |
| *Antxr1* | 1.23 | 1.17 | 4.38E-03 |
| *Adgrd1* | 1.02 | 1.17 | 4.86E-03 |
| *Col6a6* | 1.18 | 1.17 | 4.82E-03 |
| *Gpx3* | 10.21 | 1.15 | 4.97E-03 |
| *Plpp3* | 6.74 | 1.15 | 5.34E-03 |
| *Adamts2* | 1.54 | 1.14 | 6.26E-03 |
| *Efemp1* | 2.67 | 1.14 | 6.76E-03 |

**C**

| Gene | Average | Log2 FC | p-val |
|---|---|---|---|
| *Sfrp2* | 3.66 | 4.86 | 2.90E-59 |
| *Fmod* | 1.63 | 4.63 | 3.20E-35 |
| *Tnmd* | 1.37 | 4.23 | 2.60E-45 |
| *Thbs2* | 3.62 | 3.39 | 1.92E-30 |
| *Thbs4* | 18.67 | 3.11 | 9.69E-26 |
| *Igfbp3* | 2.57 | 3.01 | 1.69E-21 |
| *Col12a1* | 2.17 | 2.88 | 4.50E-21 |
| *Pdgfrl* | 4.03 | 2.75 | 8.05E-20 |
| *Cpxm2* | 1.78 | 2.66 | 7.81E-18 |
| *Mfap4* | 14.16 | 2.64 | 4.68E-18 |
| *Angptl1* | 16.18 | 2.62 | 6.04E-18 |
| *Lox* | 2.91 | 2.53 | 3.40E-16 |
| *Reck* | 1.60 | 2.48 | 1.45E-15 |
| *Itgbl1* | 2.50 | 2.43 | 6.11E-15 |
| *Eln* | 3.92 | 2.42 | 2.15E-14 |
| *C1qtnf2* | 1.66 | 2.41 | 1.29E-14 |
| *Igsf10* | 2.05 | 2.39 | 2.51E-14 |
| *Ntrk2* | 3.02 | 2.35 | 5.46E-14 |
| *Fibin* | 6.73 | 2.30 | 3.33E-13 |
| *Tmem119* | 1.13 | 2.17 | 1.44E-11 |
| *Cilp* | 16.72 | 2.16 | 1.34E-11 |
| *Tcf7l2* | 2.01 | 2.08 | 1.02E-10 |
| *Col14a1* | 10.32 | 2.06 | 1.34E-10 |
| *Sfrp1* | 3.28 | 2.05 | 3.47E-10 |
| *Mfap2* | 1.22 | 1.96 | 2.57E-09 |
| *Gas1* | 19.28 | 1.96 | 1.57E-09 |
| *Spon2* | 1.18 | 1.90 | 3.01E-08 |
| *Olfml3* | 2.81 | 1.85 | 2.37E-08 |
| *Col8a1* | 4.00 | 1.84 | 4.15E-08 |
| *Ccn2* | 5.66 | 1.84 | 6.08E-08 |
| *Ogn* | 15.44 | 1.81 | 4.42E-08 |
| *Pcsk5* | 1.37 | 1.81 | 6.81E-08 |
| *Pam* | 5.10 | 1.80 | 5.86E-08 |
| *Sfrp4* | 2.01 | 1.79 | 2.50E-06 |
| *Angptl2* | 1.03 | 1.79 | 1.15E-07 |
| *Fxyd6* | 3.54 | 1.78 | 1.04E-07 |
| *Svil* | 2.82 | 1.77 | 1.36E-07 |
| *Col1a1* | 12.58 | 1.76 | 1.59E-07 |
| *Smoc2* | 13.19 | 1.76 | 1.63E-07 |
| *Col1a2* | 30.30 | 1.73 | 2.28E-07 |
| *Svep1* | 2.62 | 1.73 | 3.22E-07 |
| *Mgp* | 31.77 | 1.72 | 4.23E-07 |
| *Cpxm1* | 2.45 | 1.70 | 6.74E-07 |
| *Rbp1* | 7.13 | 1.67 | 8.81E-07 |
| *Mmp23* | 2.96 | 1.67 | 1.04E-06 |
| *Itm2a* | 5.62 | 1.65 | 1.33E-06 |
| *Nbl1* | 10.18 | 1.63 | 2.01E-06 |
| *Aspn* | 11.67 | 1.62 | 2.53E-06 |
| *Fbln1* | 6.17 | 1.62 | 2.33E-06 |
| *C7* | 3.04 | 1.62 | 3.55E-06 |

**D**

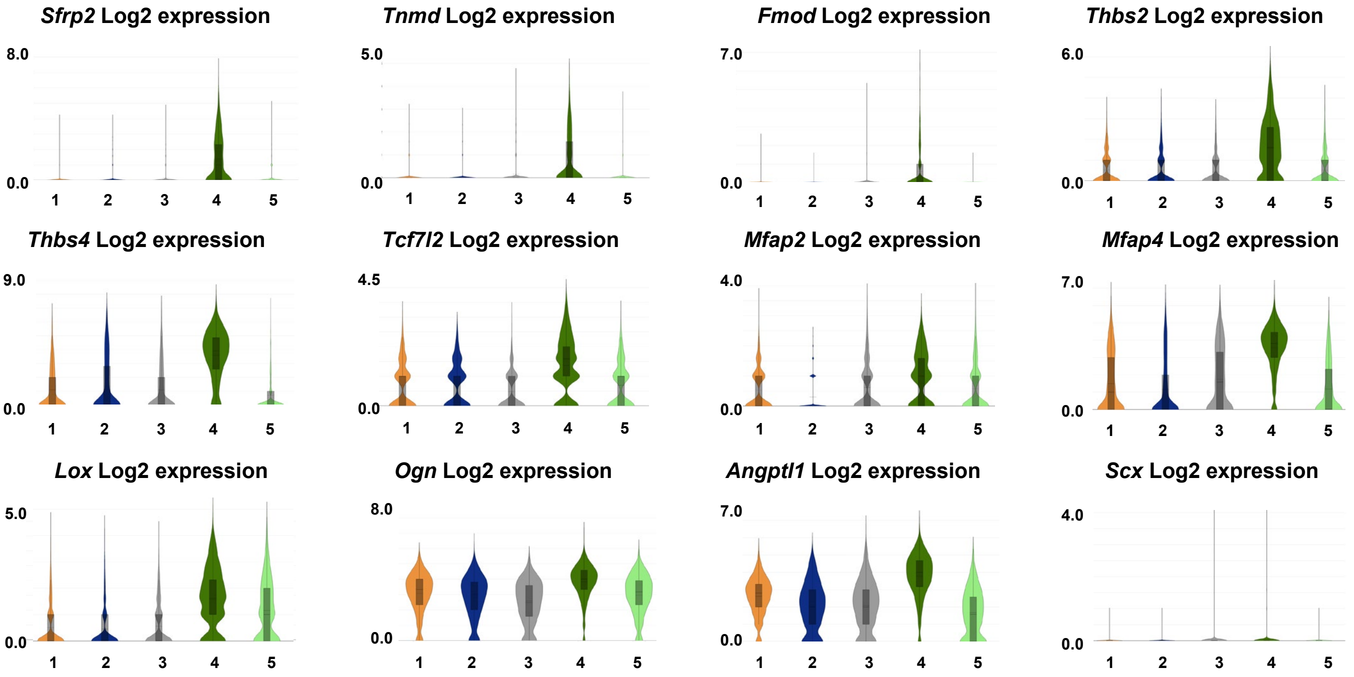


**Figure S3.** **(A)** FAP3 sub-population: Top 50 enriched genes. **(B)** FAP5 sub-population: Top 50 enriched genes. **(C)** FAP4 sub-population: Top 50 enriched genes. **(D)** Violin plots demonstrating sub-population-specific expression of selected matricellular and regulatory molecules enriched in FAP4, as well as tenocyte marker scleraixs (*Scx*). FAP1, FAP2, FAP3, FAP4 and FAP5 respectively labeled 1, 2, 3, 4, 5.

### Figure S4: Genes enriched in *Thy1*-expressing cells

**A**

| Gene | Average | Log2 FC | p-val |
|---|---|---|---|
| *Thy1* | 1.58 | 4.16 | 1.96E-58 |
| *Sema3c* | 1.98 | 2.29 | 1.52E-16 |
| *Pla1a* | 2.12 | 2.26 | 2.10E-16 |
| *Ccn3* | 1.08 | 2.24 | 2.55E-15 |
| *Cd55* | 5.43 | 2.16 | 3.56E-15 |
| *Fbn1* | 12.59 | 2.01 | 5.95E-13 |
| *Pcsk6* | 6.22 | 1.98 | 1.28E-12 |
| *Adamts5* | 4.59 | 1.76 | 7.37E-10 |
| *Adgrd1* | 1.24 | 1.73 | 2.16E-09 |
| *Uap1* | 3.11 | 1.66 | 1.35E-08 |
| *Pcolce2* | 4.09 | 1.64 | 1.84E-08 |
| *Cd248* | 4.69 | 1.63 | 2.57E-08 |
| *Dbn1* | 1.29 | 1.62 | 3.34E-08 |
| *Gfpt2* | 3.05 | 1.58 | 8.61E-08 |
| *Dpp4* | 1.06 | 1.59 | 1.08E-07 |
| *Has1* | 1.66 | 1.62 | 1.27E-07 |
| *Zfp385a* | 1.03 | 1.56 | 1.39E-07 |
| *Pi16* | 24.34 | 1.54 | 1.97E-07 |
| *Creb5* | 1.72 | 1.54 | 2.10E-07 |
| *Prss23* | 1.69 | 1.54 | 2.51E-07 |
| *Mfap5* | 20.78 | 1.52 | 2.64E-07 |
| *Ugdh* | 2.19 | 1.54 | 2.91E-07 |
| *Vcan* | 2.32 | 1.52 | 3.27E-07 |
| *Fstl1* | 12.30 | 1.49 | 4.64E-07 |
| *Ackr3* | 4.97 | 1.49 | 4.82E-07 |
| *Tnxb* | 8.74 | 1.46 | 9.36E-07 |
| *Anxa3* | 2.91 | 1.46 | 1.14E-06 |
| *Axl* | 3.75 | 1.43 | 1.94E-06 |
| *Loxl2* | 1.48 | 1.43 | 1.98E-06 |
| *Ugp2* | 2.76 | 1.42 | 2.15E-06 |
| *Fndc1* | 3.71 | 1.41 | 2.83E-06 |
| *Col5a3* | 2.16 | 1.41 | 3.16E-06 |
| *Bmp1* | 1.57 | 1.40 | 3.57E-06 |
| *Lrrn4cl* | 1.36 | 1.39 | 5.36E-06 |
| *Ccl11* | 3.65 | 1.39 | 7.36E-06 |
| *Oaf* | 3.08 | 1.36 | 8.24E-06 |
| *Myoc* | 12.15 | 1.36 | 9.14E-06 |
| *Efemp1* | 2.78 | 1.35 | 1.22E-05 |
| *Aspn* | 9.41 | 1.34 | 1.38E-05 |
| *Fap* | 1.04 | 1.32 | 1.92E-05 |
| *Nid1* | 8.41 | 1.31 | 1.97E-05 |
| *Sdc2* | 3.72 | 1.31 | 2.21E-05 |
| *Clec3b* | 14.83 | 1.30 | 2.21E-05 |
| *Cd34* | 8.35 | 1.30 | 2.51E-05 |
| *Itih5* | 4.64 | 1.28 | 3.45E-05 |
| *Lamb1* | 2.18 | 1.28 | 3.94E-05 |
| *Emilin2* | 1.81 | 1.28 | 3.95E-05 |
| *Dpt* | 12.97 | 1.27 | 4.34E-05 |
| *Tmem100* | 2.23 | 1.28 | 4.64E-05 |
| *Ddr2* | 1.90 | 1.24 | 7.45E-05 |

**B**

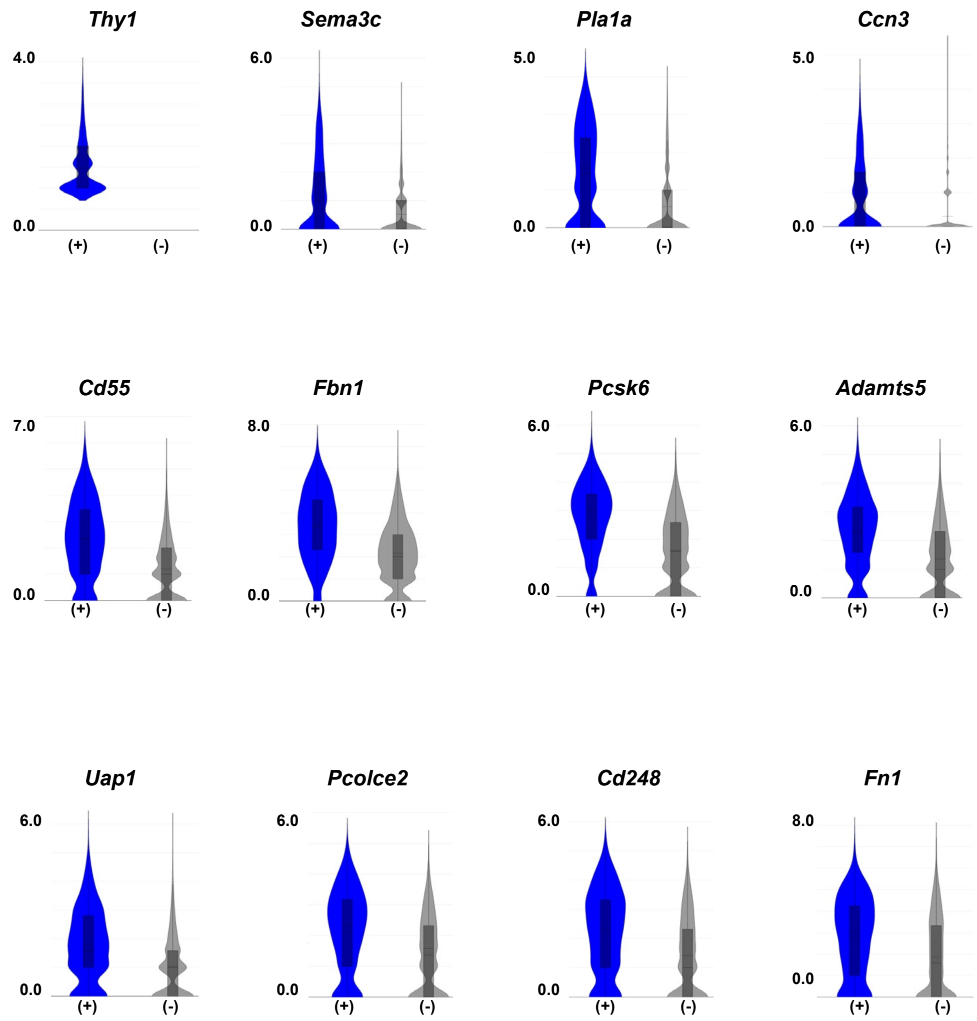


**Figure S4. (A)** *Thy-1* expressing FAPs: Top 50 enriched genes. FAP2 marker genes highlighted in blue. **(B)** Violin plots indicating expression of selected FAP2-enriched genes in *Thy1*-expressing (+) and *Thy1*-negative (-) cells.

**Figure S5: High fat diet-fed *Thbs1*$^{-/-}$ mice show similar weight gain, total body adiposity and glucose intolerance to wild type mice fed the same diet**

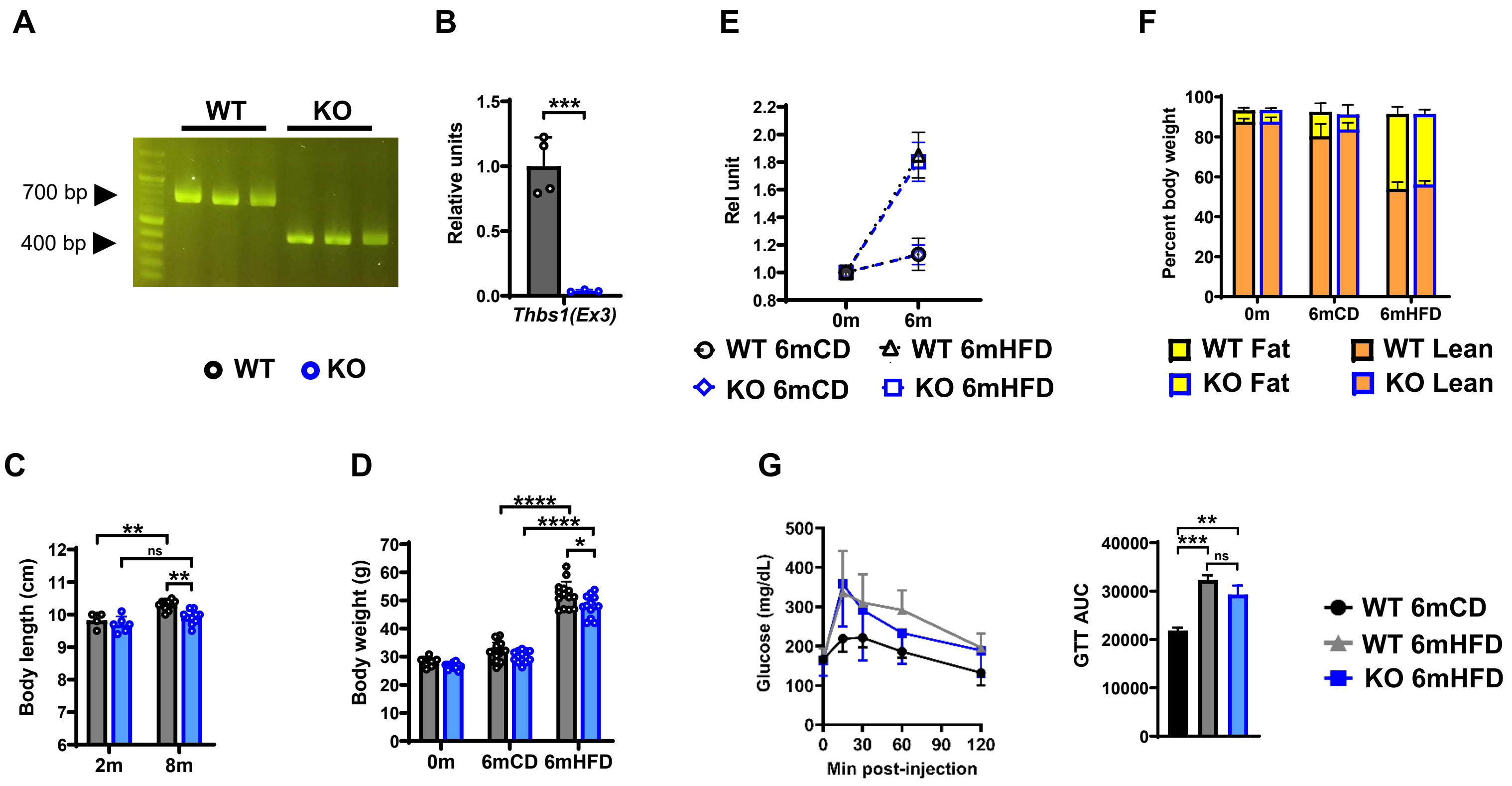


**Figure S5. (A)** Tail genotyping of *Thbs1*$^{-/-}$ (KO) mouse model. Representative gel showing PCR bands specific to wild type C57Bl/6J (WT) and KO animals. **(B)** QPCR analysis of costal diaphragm tissue from WT and KO mice using primers specifically directed against exon 3. Note that targeting vector was designed to replace exon 2, intron 3, and exon 3 of the endogenous gene with a phosphoglycerate kinase–neomycin resistance cassette (PGK-neo) cassette **[44]**. n= 4 samples per group. **(C)** Body length (nose to anus) of 2-month-old (2m) and 8-month-old (8m) WT and KO mice. n= 4-10 mice per group. **(D)** Total body weight of WT and KO mice subjected to 6-month diet time course beginning at 2-months-old (0m time point) and ending at 8-months-old (6m time point). Control diet (CD), high fat diet (HFD). n= 8-20 mice per group. **(E)** Weight gain (normalized to baseline) in CD and HFD-fed WT and KO mice. n= 8-20 mice per group. **(F)** NMR-based body composition of WT and KO mice at baseline (2-months-old, 0m time point) and following 6-month CD or HFD feeding. n= 6-8 mice per group. **(G)** Intraperitoneal glucose tolerance test performed on 6m HFD-fed WT and KO mice; as well as age matched WT mice fed CD. Line graph indicates blood glucose values at baseline (0) and 15, 30, 60, 90 and 120 minutes (min) following intraperitoneal injection of 1 mg/kg dextrose. Bar graph compares area under the curve (AUC) for each line. n= 6-8 mice per group. Error bars indicate SDM. $^*p<0.05$, $^{**}p<0.01$, $^{***}p<0.001$, $^{****}p<0.001$.

**Figure S6: *Thbs1* ablation has minimal effect on organ and muscle weights in HFD-fed mice**

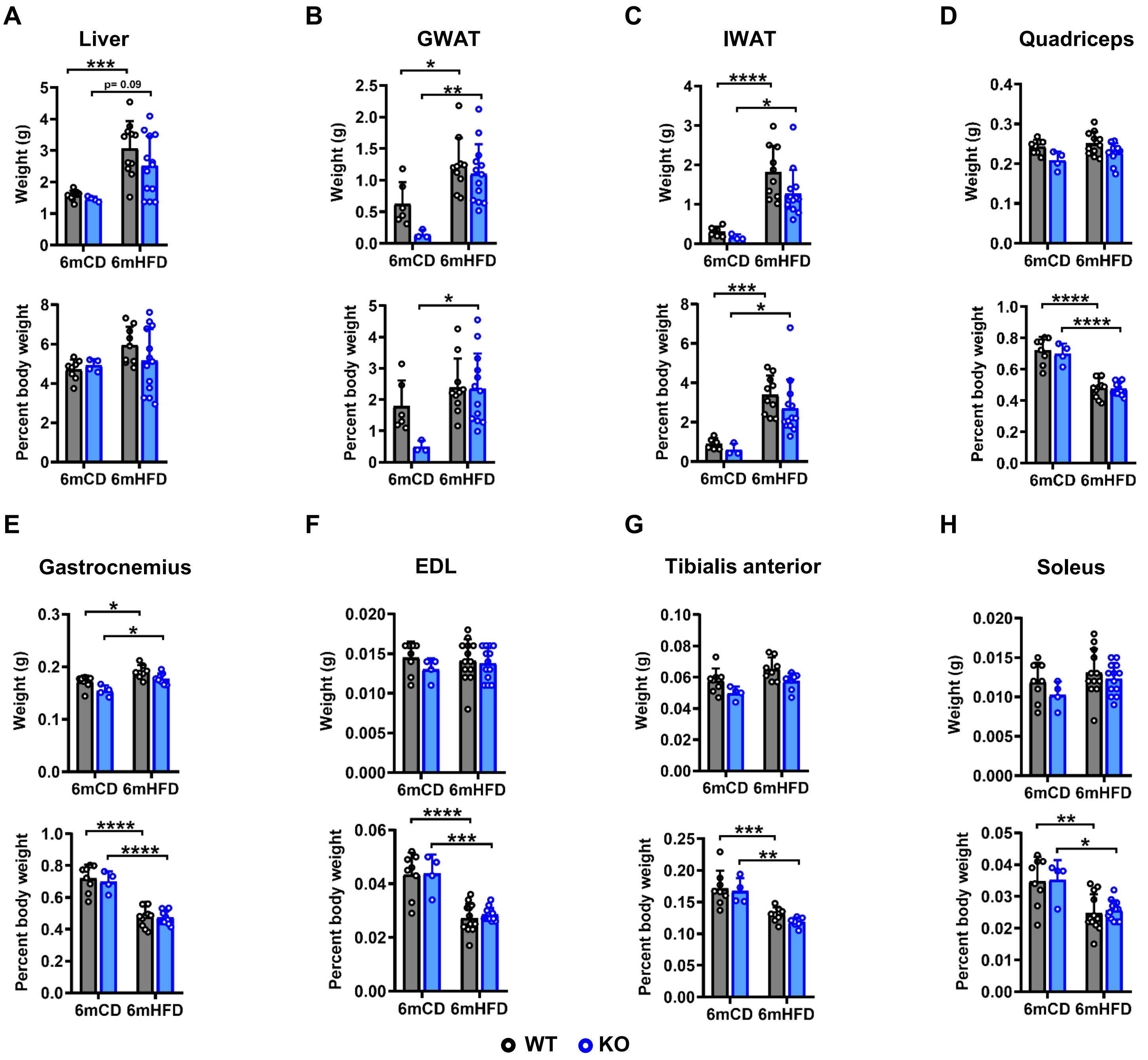


**Figure S6.** Weights (absolute, top panels; % total body weight, bottom panels) of **(A)** Liver, **(B)** Gonadal white adipose tissue (GWAT), **(C)** Inguinal white adipose tissue (IWAT), **(D)** Quadriceps, **(E)** Gastrocnemius, **(F)** Extensor digitorum longus (EDL), **(G)** Tibialis anterior, and **(H)** Soleus. All isolated from WT and KO mice after 6-month control diet (6mCD) or 6-month high fat diet (6mHFD) feeding. Fat depot weights represent combined weight of bilateral depots. Muscle weights indicate weight of a single muscle. n= 4-15 animals per group. Error bars indicate SDM. *p<0.05, **p<0.01, ***p<0.001, ****p<0.0001.

#### Figure S7: Inter-group comparisons: Differentially expressed genes

**A**

*Thy1*-expressing cell marker genes

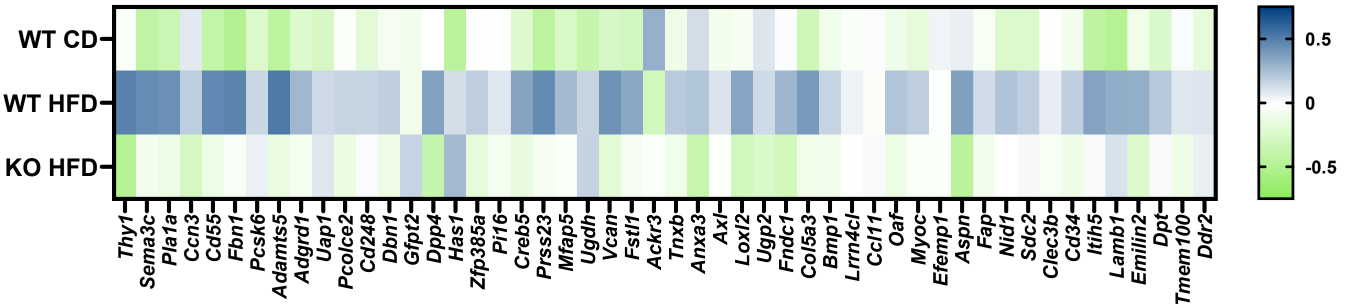


**B**

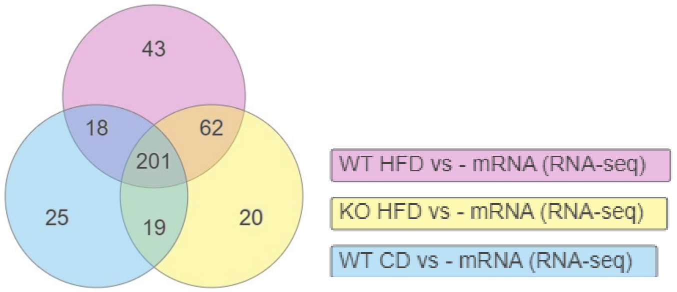


**C**

##### Common to WT HFD and KO HFD

| Gene | WT HFD | pval | KO HFD | pval | WT CD | pval |
|---|---|---|---|---|---|---|
| Fbn1 | 1.392 | 1.20E-06 | 0.956 | 0.002 | 0.452 | 0.371 |
| Adamts5 | 1.357 | 2.48E-06 | 0.773 | 0.019 | 0.452 | 0.37 |
| Sema3c | 1.341 | 6.48E-06 | 0.9 | 0.005 | 0.514 | 0.276 |
| Itih5 | 1.286 | 1.06E-05 | 0.977 | 0.001 | 0.502 | 0.275 |
| Dclk1 | 1.279 | 1.43E-05 | 0.983 | 0.001 | 0.539 | 0.222 |
| Cd55 | 1.261 | 2.03E-05 | 0.75 | 0.025 | 0.47 | 0.338 |
| Col6a3 | 1.249 | 2.25E-05 | 1.021 | 6.31E-04 | 0.538 | 0.217 |
| Col5a3 | 1.25 | 2.34E-05 | 0.818 | 0.011 | 0.541 | 0.215 |
| Col3a1 | 1.246 | 2.37E-05 | 0.87 | 0.006 | 0.583 | 0.159 |
| Lamb1 | 1.173 | 9.81E-05 | 1.006 | 8.28E-04 | 0.392 | 0.49 |
| Ccdc80 | 1.164 | 1.08E-04 | 0.957 | 0.002 | 0.563 | 0.183 |
| Sparc | 1.119 | 2.19E-04 | 0.769 | 0.019 | 0.512 | 0.254 |
| Ahnak2 | 1.088 | 4.21E-04 | 0.641 | 0.074 | 0.512 | 0.261 |
| Golim4 | 1.073 | 5.13E-04 | 0.623 | 0.086 | 0.514 | 0.255 |
| Igsf10 | 1.072 | 6.02E-04 | 1.225 | 1.95E-05 | 0.532 | 0.235 |
| Ly6a | 1.034 | 9.03E-04 | 0.642 | 0.071 | 0.283 | 0.696 |
| Lama4 | 1.026 | 0.001 | 0.78 | 0.018 | 0.372 | 0.534 |
| Creb5 | 1.016 | 0.001 | 0.589 | 0.116 | 0.466 | 0.345 |
| Has1 | 1.038 | 0.002 | 1.25 | 2.53E-05 | 0.477 | 0.357 |
| Steap3 | 0.996 | 0.002 | 1.278 | 5.29E-06 | 0.385 | 0.503 |
| Egfr | 0.989 | 0.002 | 0.792 | 0.015 | 0.45 | 0.373 |
| Amotl1 | 0.98 | 0.002 | 0.838 | 0.009 | 0.512 | 0.259 |
| Ppic | 0.98 | 0.002 | 0.815 | 0.012 | 0.445 | 0.384 |
| Csf1 | 0.968 | 0.003 | 0.983 | 0.001 | 0.556 | 0.195 |
| Ugdh | 0.971 | 0.003 | 1.036 | 6.18E-04 | 0.463 | 0.36 |
| Prss23 | 0.971 | 0.003 | 0.597 | 0.114 | 0.184 | 0.887 |
| Uap1 | 0.963 | 0.003 | 0.935 | 0.003 | 0.565 | 0.186 |
| Fbln5 | 0.948 | 0.003 | 0.823 | 0.01 | 0.376 | 0.521 |
| Hspg2 | 0.93 | 0.004 | 0.758 | 0.022 | 0.563 | 0.183 |
| Col4a1 | 0.926 | 0.004 | 0.69 | 0.044 | 0.183 | 0.88 |
| Antxr2 | 0.924 | 0.004 | 0.585 | 0.117 | 0.434 | 0.398 |
| Ifi205 | 0.917 | 0.005 | 1.132 | 1.02E-04 | 0.531 | 0.231 |
| Sntb2 | 0.882 | 0.007 | 0.716 | 0.034 | 0.399 | 0.472 |
| Frmd6 | 0.843 | 0.012 | 0.702 | 0.04 | 0.481 | 0.313 |
| Col4a2 | 0.832 | 0.014 | 0.598 | 0.108 | 0.154 | 0.939 |
| Nfix | 0.818 | 0.015 | 0.757 | 0.021 | 0.568 | 0.174 |
| Akap12 | 0.821 | 0.016 | 0.698 | 0.044 | 0.472 | 0.333 |
| Ablim1 | 0.806 | 0.018 | 0.629 | 0.08 | 0.572 | 0.173 |
| Agap1 | 0.805 | 0.019 | 0.676 | 0.053 | 0.514 | 0.258 |
| Cavin3 | 0.792 | 0.021 | 0.657 | 0.061 | 0.457 | 0.355 |
| Serpinb6a | 0.791 | 0.021 | 0.735 | 0.027 | 0.499 | 0.274 |
| Timp2 | 0.78 | 0.023 | 0.683 | 0.047 | 0.543 | 0.205 |
| Vat1 | 0.784 | 0.023 | 0.688 | 0.046 | 0.495 | 0.287 |
| Ifi211 | 0.783 | 0.024 | 1.045 | 4.59E-04 | 0.438 | 0.395 |
| Ebf1 | 0.776 | 0.025 | 0.638 | 0.073 | 0.477 | 0.316 |
| Colec12 | 0.76 | 0.03 | 0.801 | 0.014 | 0.542 | 0.213 |
| Nav1 | 0.744 | 0.035 | 0.826 | 0.01 | 0.49 | 0.292 |
| Cyth3 | 0.742 | 0.037 | 0.732 | 0.03 | 0.484 | 0.309 |
| Sash1 | 0.736 | 0.038 | 0.683 | 0.048 | 0.158 | 0.929 |
| Nedd4 | 0.72 | 0.043 | 0.716 | 0.033 | 0.392 | 0.482 |
| Itm2a | 0.699 | 0.055 | 0.699 | 0.042 | 0.448 | 0.375 |
| Zbtb20 | 0.692 | 0.056 | 0.674 | 0.051 | 0.35 | 0.557 |
| Lima1 | 0.681 | 0.064 | 0.594 | 0.108 | 0.437 | 0.393 |
| Nfia | 0.678 | 0.065 | 0.68 | 0.049 | 0.468 | 0.332 |
| C3 | 0.676 | 0.069 | 1.061 | 3.40E-04 | 0.526 | 0.237 |
| Ifi204 | 0.668 | 0.076 | 0.885 | 0.005 | 0.236 | 0.79 |
| Nfib | 0.644 | 0.086 | 0.739 | 0.026 | 0.41 | 0.444 |
| Thbs1 | 0.658 | 0.097 | 0.601 | 0.123 | 0.361 | 0.559 |
| Man1a | 0.593 | 0.131 | 0.618 | 0.087 | 0.486 | 0.298 |
| Bmp4 | 0.598 | 0.133 | 0.78 | 0.019 | 0.536 | 0.228 |
| Tspan3 | 0.576 | 0.15 | 0.608 | 0.096 | 0.508 | 0.262 |
| Fcgrt | 0.573 | 0.151 | 0.673 | 0.052 | 0.539 | 0.213 |
| Ecm1 | 0.397 | 0.427 | 0.631 | 0.078 | 0.387 | 0.497 |
| Errfi1 | 0.389 | 0.444 | 0.778 | 0.017 | 0.142 | 0.953 |

**D**

##### WT HFD-specific

| Gene | WT HFD | pval | KO HFD | pval | WT CD | pval |
|---|---|---|---|---|---|---|
| Zeb1 | 0.881 | 0.008 | 0.566 | 0.138 | 0.491 | 0.297 |
| Tgfbr2 | 0.863 | 0.009 | 0.517 | 0.194 | 0.437 | 0.393 |
| Prg4 | 0.892 | 0.011 | 0.227 | 0.767 | 0.072 | 1 |
| Igf2r | 0.853 | 0.011 | 0.555 | 0.149 | 0.402 | 0.468 |
| Emilin2 | 0.845 | 0.012 | 0.392 | 0.419 | 0.461 | 0.356 |
| Cald1 | 0.805 | 0.018 | 0.569 | 0.132 | 0.322 | 0.618 |
| Marcks | 0.755 | 0.03 | 0.508 | 0.204 | -0.058 | 1 |
| P4ha1 | 0.757 | 0.031 | 0.53 | 0.179 | 0.489 | 0.297 |
| Arl4a | 0.752 | 0.032 | 0.571 | 0.131 | 0.548 | 0.202 |
| Enpp2 | 0.759 | 0.035 | 0.462 | 0.294 | 0.578 | 0.178 |
| S100a16 | 0.727 | 0.042 | 0.463 | 0.277 | 0.425 | 0.415 |
| Sec31a | 0.726 | 0.042 | 0.508 | 0.207 | 0.486 | 0.302 |
| Mast4 | 0.705 | 0.053 | 0.567 | 0.137 | 0.396 | 0.48 |
| Rhoj | 0.685 | 0.061 | 0.566 | 0.135 | 0.437 | 0.392 |
| Thra | 0.686 | 0.063 | 0.423 | 0.355 | 0.507 | 0.267 |
| Glg1 | 0.672 | 0.07 | 0.465 | 0.276 | 0.223 | 0.809 |
| Nfic | 0.649 | 0.083 | 0.563 | 0.138 | 0.509 | 0.26 |
| Fermt2 | 0.646 | 0.085 | 0.46 | 0.281 | 0.417 | 0.432 |
| Dap | 0.645 | 0.087 | 0.501 | 0.216 | 0.449 | 0.371 |
| Hdlbp | 0.631 | 0.097 | 0.392 | 0.41 | 0.321 | 0.617 |
| Sar1a | 0.626 | 0.101 | 0.421 | 0.355 | 0.429 | 0.408 |
| Sptbn1 | 0.626 | 0.101 | 0.341 | 0.525 | 0.298 | 0.664 |
| Rhoc | 0.624 | 0.103 | 0.483 | 0.244 | 0.469 | 0.332 |
| Wls | 0.625 | 0.104 | 0.519 | 0.193 | 0.498 | 0.283 |
| Dnajc3 | 0.619 | 0.107 | 0.266 | 0.672 | 0.567 | 0.175 |
| Arl1 | 0.615 | 0.11 | 0.405 | 0.387 | 0.406 | 0.454 |
| Tcf4 | 0.612 | 0.112 | 0.472 | 0.26 | 0.33 | 0.595 |
| Ski | 0.611 | 0.114 | 0.475 | 0.256 | 0.4 | 0.468 |
| Lman1 | 0.61 | 0.115 | 0.334 | 0.54 | 0.361 | 0.547 |
| Anxa1 | 0.607 | 0.117 | 0.521 | 0.187 | 0.337 | 0.58 |
| Ramp2 | 0.603 | 0.123 | 0.538 | 0.169 | 0.282 | 0.701 |
| Dpysl2 | 0.599 | 0.125 | 0.444 | 0.31 | 0.52 | 0.242 |
| Copb2 | 0.599 | 0.125 | 0.43 | 0.34 | 0.217 | 0.823 |
| P4hb | 0.597 | 0.126 | 0.529 | 0.176 | 0.469 | 0.329 |
| Ly6c1 | 0.601 | 0.131 | 0.523 | 0.194 | -0.064 | 1 |
| Add3 | 0.588 | 0.135 | 0.447 | 0.304 | 0.467 | 0.336 |
| Dst | 0.587 | 0.141 | 0.336 | 0.541 | 0.154 | 0.939 |

**E**

##### KO HFD-specific

| Gene | WT HFD | pval | KO HFD | pval | WT CD | pval |
|---|---|---|---|---|---|---|
| Col6a5 | 0.256 | 0.726 | 1.882 | 1.00E-06 | 0.359 | 0.573 |
| Phlda1 | 0.324 | 0.567 | 1.005 | 8.11E-04 | 0.577 | 0.167 |
| Cxcl1 | 0.203 | 0.811 | 1.002 | 0.001 | 0.313 | 0.642 |
| Errfi1 | 0.389 | 0.444 | 0.778 | 0.017 | 0.142 | 0.953 |
| Adamts1 | 0.209 | 0.794 | 0.717 | 0.035 | 0.374 | 0.526 |
| Myc | 0.044 | 1 | 0.711 | 0.04 | 0.169 | 0.913 |
| Tent5a | 0.316 | 0.588 | 0.683 | 0.05 | 0.395 | 0.484 |
| Pnp | 0.229 | 0.749 | 0.676 | 0.05 | 0.528 | 0.229 |
| Gem | 0.267 | 0.686 | 0.678 | 0.051 | 0.43 | 0.408 |
| Fcgrt | 0.573 | 0.151 | 0.673 | 0.052 | 0.539 | 0.213 |
| Nr4a1 | -0.126 | 0.988 | 0.661 | 0.06 | 0.424 | 0.419 |
| Ifi203 | -0.006 | 1 | 0.655 | 0.066 | -0.045 | 1 |
| Ecm1 | 0.397 | 0.427 | 0.631 | 0.078 | 0.387 | 0.497 |
| Klf4 | 0.269 | 0.678 | 0.61 | 0.093 | 0.389 | 0.49 |
| Bag3 | 0.148 | 0.911 | 0.614 | 0.093 | 0.432 | 0.407 |
| Tspan3 | 0.576 | 0.15 | 0.608 | 0.096 | 0.508 | 0.262 |
| Egr1 | 0.115 | 0.964 | 0.597 | 0.105 | 0.444 | 0.378 |
| Fos | 0.293 | 0.623 | 0.591 | 0.109 | 0.332 | 0.59 |
| Cebpd | 0.191 | 0.825 | 0.591 | 0.111 | 0.077 | 1 |

**Figure S7. (A)** Heat map indicating the expression of *Thy1*-expressing cell marker genes in WT CD, WT HFD and KO HFD groups. **(B)** Venn diagram showing differentially expressed genes in diaphragm FAPs from WT CD, WT HFD, KO HFD mice. **(C)** Transcripts enriched in WT HFD and KO HFD (n= 62; orange section of Venn diagram) . **(D)** Genes uniquely enriched in WT HFD (n= 43; pink section of Venn diagram). **(E)** Genes uniquely enriched in KO HFD (n= 20; yellow section of Venn diagram). Indicated p-values are corrected for false discovery rate (FDR).

### Figure S8: Inter-group comparisons: Differentially represented gene ontology terms

**A**

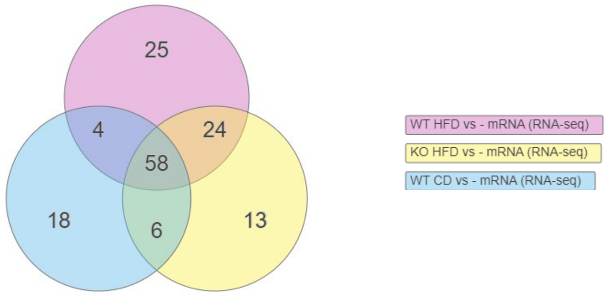


**B**

##### Common to all groups

| Biological Process | WT CD pval | WT HFD pval | KO HFD pval |
|---|---|---|---|
| External encapsulating structure organization | 1.51E-09 | 1.06E-15 | 1.69E-17 |
| Extracellular structure organization | 1.51E-09 | 1.06E-15 | 1.69E-17 |
| Extracellular matrix organization | 1.51E-09 | 1.06E-15 | 1.69E-17 |
| System development | 3.04E-06 | 4.90E-10 | 3.04E-08 |
| Anatomical structure development | 2.94E-06 | 2.62E-09 | 2.87E-07 |
| Anatomical structure morphogenesis | 3.35E-04 | 1.76E-08 | 4.37E-06 |
| Multicellular organism development | 1.01E-05 | 2.37E-08 | 3.37E-07 |
| Skeletal system development | 2.26E-04 | 7.54E-08 | 4.05E-08 |
| Multicellular organismal process | 3.73E-06 | 1.01E-07 | 2.17E-07 |
| Developmental process | 1.28E-05 | 1.51E-07 | 4.40E-06 |
| Animal organ development | 2.83E-05 | 2.56E-07 | 8.63E-07 |
| Cell adhesion | 1.58E-06 | 7.96E-07 | 3.61E-07 |
| Biological adhesion | 1.90E-06 | 1.16E-06 | 5.06E-07 |
| Connective tissue development | 2.10E-02 | 1.29E-06 | 8.71E-07 |
| Cartilage development | 3.00E-02 | 1.98E-06 | 2.14E-05 |
| Animal organ morphogenesis | 5.92E-04 | 2.33E-06 | 2.44E-05 |
| Collagen fibril organization | 3.83E-05 | 2.34E-06 | 1.01E-05 |
| Cell migration | 1.28E-05 | 2.74E-06 | 6.33E-08 |
| Tissue development | 3.35E-04 | 9.26E-06 | 1.44E-06 |
| Circulatory system development | 2.00E-03 | 1.38E-05 | 3.50E-07 |
| Localization of cell | 4.53E-05 | 1.57E-05 | 3.27E-07 |
| Cell motility | 4.53E-05 | 1.57E-05 | 3.27E-07 |
| Enzyme linked receptor protein signaling pathway | 1.25E-04 | 1.57E-05 | 8.29E-04 |
| Locomotion | 4.81E-05 | 1.57E-05 | 2.87E-07 |
| Tube development | 3.00E-03 | 1.66E-05 | 7.38E-06 |
| Vasculature development | 4.00E-03 | 1.77E-05 | 8.54E-07 |
| Blood vessel development | 7.00E-03 | 2.87E-05 | 4.10E-06 |
| Tube morphogenesis | 2.60E-02 | 3.64E-05 | 2.62E-05 |
| Cellular response to growth factor stimulus | 1.90E-02 | 4.69E-05 | 2.00E-03 |
| Movement of cell or subcellular component | 6.95E-04 | 1.25E-04 | 1.67E-06 |
| Response to growth factor | 4.80E-02 | 1.40E-04 | 8.00E-03 |
| Cell differentiation | 2.26E-04 | 1.92E-04 | 2.62E-05 |
| Cell-substrate adhesion | 5.00E-03 | 4.24E-04 | 2.53E-04 |
| Cellular developmental process | 4.71E-04 | 5.78E-04 | 8.77E-05 |
| Cell surface receptor signaling pathway | 1.56E-04 | 6.83E-04 | 1.96E-04 |
| Regulation of multicellular organismal process | 1.40E-02 | 6.83E-04 | 1.00E-03 |
| Supramolecular fiber organization | 2.00E-03 | 1.00E-03 | 2.00E-02 |
| Ossification | 3.60E-02 | 2.00E-03 | 7.00E-03 |
| Regulation of developmental process | 2.30E-02 | 2.00E-03 | 1.30E-02 |
| Regulation of BMP signaling pathway | 2.00E-03 | 2.00E-03 | 6.00E-03 |
| Biomineral tissue development | 2.00E-03 | 2.00E-03 | 1.00E-03 |
| Biomineralization | 2.00E-03 | 2.00E-03 | 1.00E-03 |
| Wound healing | 1.60E-02 | 5.00E-03 | 1.00E-02 |
| Biological regulation | 2.90E-02 | 5.00E-03 | 1.00E-02 |
| Collagen metabolic process | 4.15E-04 | 5.00E-03 | 3.87E-04 |
| Response to wounding | 7.00E-03 | 5.00E-03 | 3.00E-03 |
| Response to BMP | 4.00E-03 | 6.00E-03 | 2.00E-03 |
| Cellular response to BMP stimulus | 4.00E-03 | 6.00E-03 | 2.00E-03 |
| BMP signaling pathway | 4.00E-03 | 6.00E-03 | 2.00E-03 |
| Transmembrane receptor protein tyrosine kinase signaling pathway | 3.00E-03 | 7.00E-03 | 4.00E-03 |
| Blood vessel remodeling | 4.80E-02 | 1.30E-02 | 7.00E-03 |
| Bone mineralization | 2.00E-03 | 1.30E-02 | 1.00E-03 |
| Regulation of cell motility | 4.40E-02 | 1.60E-02 | 2.00E-03 |
| Regulation of locomotion | 4.90E-02 | 2.00E-02 | 2.00E-03 |
| Regulation of response to stimulus | 4.00E-03 | 2.40E-02 | 5.00E-03 |
| Negative regulation of multicellular organismal process | 4.00E-02 | 2.40E-02 | 7.00E-03 |
| System process | 6.95E-04 | 3.60E-02 | 6.00E-03 |
| Negative regulation of cell population proliferation | 2.80E-04 | 3.60E-02 | 9.00E-03 |

**C**

##### Specific to WT CD

| Biological Process | P-val (corr for FDR) |
|---|---|
| Negative regulation of signal transduction | 0.0080 |
| Hormone metabolic process | 0.0100 |
| Negative regulation of cell communication | 0.0120 |
| Negative regulation of signaling | 0.0120 |
| Regulation of blood pressure | 0.0140 |
| Regulation of systemic arterial blood pressure | 0.0180 |
| Sensory perception | 0.0210 |
| Circulatory system process | 0.0210 |
| Blood vessel diameter maintenance | 0.0290 |
| Regulation of tube size | 0.0290 |
| Regulation of tube diameter | 0.0290 |
| Cell development | 0.0290 |
| Regulation of hormone levels | 0.0360 |
| Regulation of signal transduction | 0.0400 |
| Regulation of tissue remodeling | 0.0480 |
| Regulation of response to external stimulus | 0.0480 |
| Regulation of insulin-like growth factor receptor signaling pathway | 0.0480 |

**D**

##### Specific to KO HFD

| Biological Process | P-val (corr for FDR) |
|---|---|
| Renal system development | 0.0020 |
| Regulation of ossification | 0.0050 |
| Regulation of epithelial cell proliferation | 0.0060 |
| Kidney development | 0.0060 |
| Nephron development | 0.0150 |
| Regulation of cell population proliferation | 0.0170 |
| Negative regulation of peptidase activity | 0.0170 |
| Negative regulation of endopeptidase activity | 0.0250 |
| Angiogenesis | 0.0250 |
| Cell population proliferation | 0.0270 |
| Regulation of bone mineralization | 0.0320 |
| Regulation of phosphatidylinositol 3-kinase signaling | 0.0350 |
| Glomerulus development | 0.0350 |

**E**

##### Common to WT HFD and KO HFD

| Biological Process | P-val (corr for FDR) |
|---|---|
| Bone development | 6.80E-04 |
| Skeletal system morphogenesis | 7.00E-03 |
| Extracellular matrix assembly | 2.00E-03 |
| Negative regulation of cellular response to growth factor stimulus | 3.50E-02 |
| Blood vessel morphogenesis | 2.00E-03 |
| Chondrocyte differentiation | 1.40E-02 |
| Regulation of multicellular organismal development | 1.40E-02 |
| Heart development | 2.00E-03 |
| Complement activation | 5.00E-03 |
| Anatomical structure formation involved in morphogenesis | 1.50E-02 |
| Complement activation, classical pathway | 7.00E-03 |
| Response to stimulus | 1.00E-02 |
| Urogenital system development | 3.87E-04 |
| Regulation of biological process | 3.80E-02 |
| Epithelium development | 1.10E-02 |
| Regulation of cell migration | 3.00E-03 |
| Reproductive structure development | 1.90E-02 |
| Reproductive system development | 1.90E-02 |
| Response to chemical | 1.70E-02 |
| Regulation of cellular component movement | 4.00E-03 |
| Prostate gland morphogenesis | 2.10E-02 |
| Prostate gland epithelium morphogenesis | 2.10E-02 |
| Elastic fiber assembly | 2.10E-02 |
| Humoral immune response mediated by circulating immunoglobulin | 3.20E-02 |

**Figure S8. (A)** Venn diagram showing differentially enriched gene ontology terms (biological processes) in WT CD, WT HFD and KO HFD diaphragm FAPs. **(B)** Biological processes common to all groups (n= 58). **(C)** Biological processes enriched in WT CD diaphragm FAPs (n= 18; blue section of Venn diagram). **(D)** Biological processes enriched in KO HFD diaphragm FAPs (n= 13; yellow section of Venn diagram). **(E)** Biological processes common to WT HFD and KO HFD FAPs (n= 24; orange section of Venn diagram). Indicated p-values are corrected for false discovery rate (FDR).

### Figure S9: TGFβ, PDGF and THBS1 receptor expression in FAP sub-populations

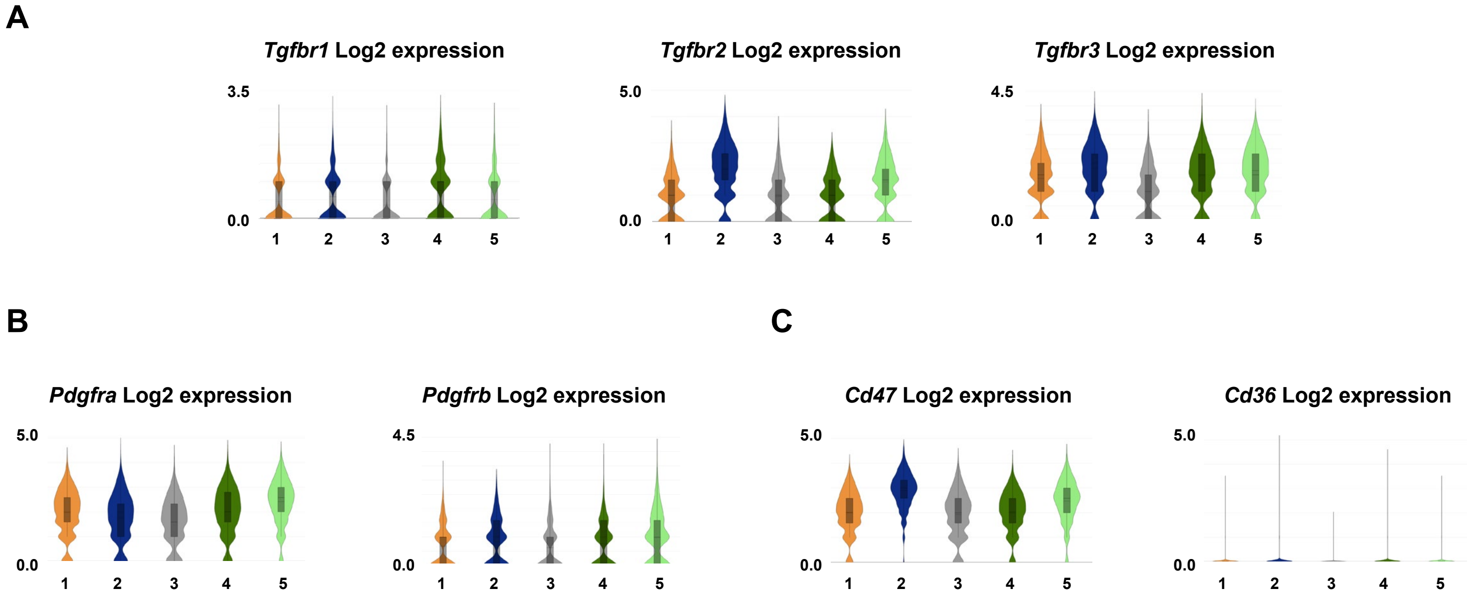


**Figure S9. (A-C)** Expression of TGFβ receptors (*Tgfbr1*, *Tgfbr2*, *Tgfbr3*); PDGF receptors (*Pdgfra*, *Pdgfrb*) and THBS1 receptors (*Cd36*, *Cd47*) in individual FAP sub-populations. FAP1, FAP2, FAP3, FAP4 and FAP5 respectively labeled 1, 2, 3, 4, 5.

#### Figure S10: Effect of DIO and *Thbs1* ablation on diaphragm macrophages

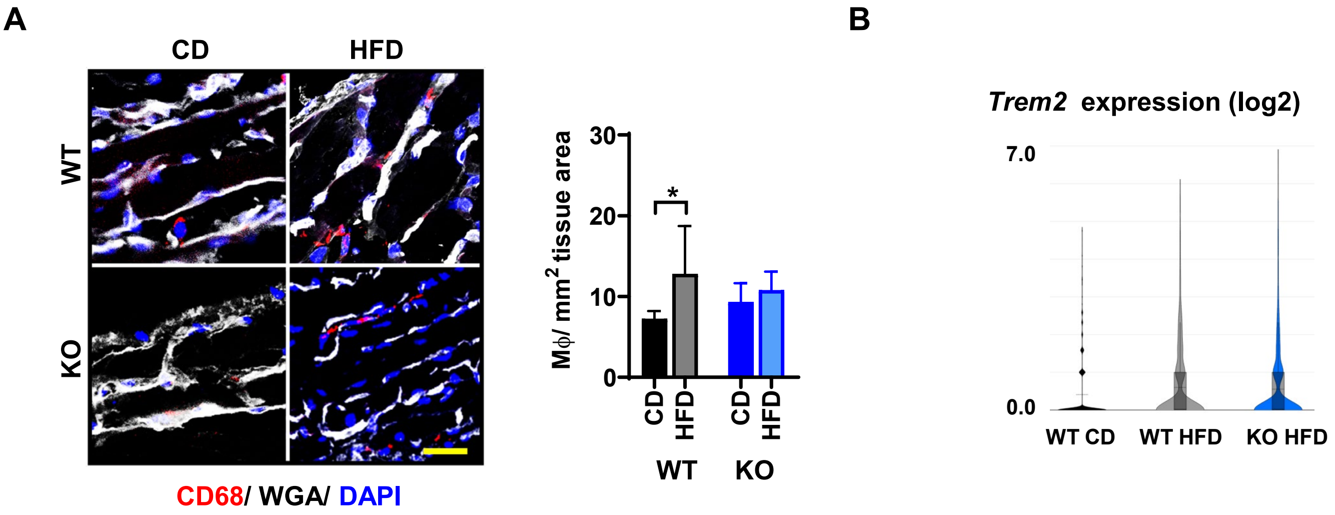


**Figure S10. (A)** CD68 immunohistochemistry in costal diaphragm longitudinal sections from wild type (WT) and *Thbs1*$^{-/-}$ (KO) mice fed either control diet (CD) or high fat diet (HFD) for 6 months. Wheat germ agglutinin, WGA. Scale 20 µm. Bar graph shows number of CD68 immunopositive cells per $mm^2$ cross sectional area. n= 4-5 per group. **(B)** Expression of obesity-associated macrophage-specific *Trem2* in macrophage sub-clusters and (WT CD, WT HFD and KO HFD) groups. Error bars indicate SDM. *p<0.05.

**Figure S11: Diaphragm histology and physiology parameters**

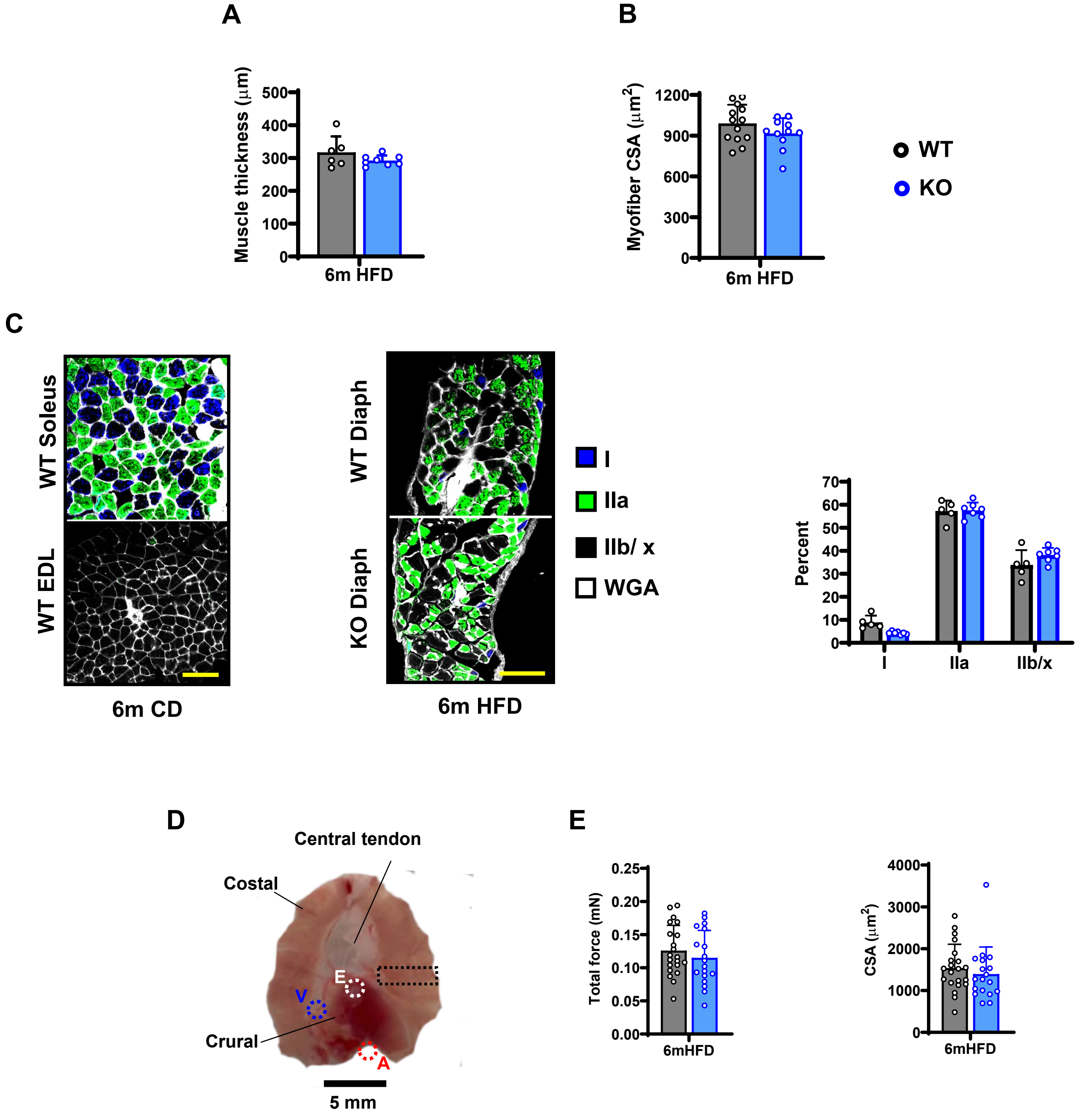


**Figure S11. (A)** Diaphragm thickness measured on FFPE longitudinal sections. 5-7 animals per group, 3 non-consecutive sections per animal, 10-15 thickness measurements per section. **(B)** Myofiber size quantification (fiber cross-sectional area) 5-10 images of 3 non-consecutive transverse sections from 4 animals per group. **(C)** Transverse sections from unfixed frozen samples of extensor digitorum longus (EDL) and soleus stained with fiber typing antibodies—positive controls for fiber typing antibodies. Type I (slow oxidative), type IIa (fast oxidative), type IIb/ x (fast glycolytic). Transverse sections from unfixed frozen samples of diaphragm stained with fiber typing antibodies. Bar graph shows percentage of each fiber type in diaphragm samples from each group. n= 4-8 animals/ group. **(D)** Diagram of mouse diaphragm indicating costal muscle, crural muscle and central tendon region, as well as positions of aorta (A), descending vena cava (V) and esophagus (E). B lack dashed line indicates position of strips isolated for isometric force testing. **(E)** Isometric force measurement performed on single myofibers isolated from 6mHFD WT and KO mice (n= 4-5 animals per group; 5 fibers per animal). Graphs indicate total isometric force and fiber cross sectional area. Error bars indicate SDM.

### Table S1: Antibodies

| | Primary antibodies | | Secondary antibodies | |
|---|---|---|---|---|
| Experiment | Target, animal, conjugate, company, catalog # | Dilution | Target, animal, conjugate, company, catalog # | Dilution |
| Cell culture ICC | Ki67, rabbit, unconjugated Abcam, ab15580 | 1:250 | Goat anti-rabbit, Alexa Fluor 594, Invitrogen, A11072 | 1:250 |
| | Fibronectin, rabbit, unconjugated, Sigma Aldrich F3648 | 1:250 | Goat anti-rabbit, Alexa Fluor 594, Invitrogen, A11072 | 1:250 |
| Tissue (frozen secions) IHC | PDGFRα, goat, unconjugated R&D Systems, AF1062 | 1:400 | Donkey anti-goat, Alexa Fluor 594, Invitrogen, A-32758 | 1:2000 |
| | CD68, rat, unconjugated, BioRad, MCA1957 | 1:200 | Donkey anti-rat, Alexa Fluor 594, Invitrogen, A-21209 | 1:200 |
| | Myosin heavy chain Type I, mouse, DHSB, BA-D5-c | 1:300 | Goat anti-mouse, Alexa Fluor 647, Invitrogen, A-21242 | 1:300 |
| | Myosin heavy chain Type IIA, mouse, DHSB, SC-71-c | 1:300 | Goat anti-mouse, Alexa Fluor 488, Invitrogen, A-21121 | 1:300 |
| Tissue (FFPE sections) IHC | Perilipin-1, rabbit, unconjugated, Cell Signaling, 3470S | 1:250 | Goat anti-rabbit, Alexa Fluor 594, Invitrogen, A-11072 | 1:1000 |
| | THY1/ CD90, rabbit, unconjugated, Bioss, BS-0778R | 1:400 | Goat anti-rabbit, Alexa Fluor 594, Invitrogen, A-11072 | 1:1000 |
| | Fibronectin, rabbit, unconjugated, Sigma Aldrich F3648 | 1:400 | Goat anti-rabbit, Alexa Fluor 594, Invitrogen, A-11072 | 1:1000 |
| Flow cytometry | CD31, rat, PE, eBioscience, 12-0311-81 | 1:1000 | N/A | N/A |
| | CD45, rat, PE, eBioscience, 12-0451-82 | 1:1000 | N/A | N/A |
| | Integrin α7, rat, PE, Invitrogen, MA5-23608 | 1:1000 | N/A | N/A |
| | Sca-1, rat, APC, eBioschence, 17-5981-82 | 1:1000 | N/A | N/A |

**Table S1.** ICC, immunocytochemistry. IHC, immunohistochemistry. FFPE, formalin-fixed, paraffin-embedded. DHSB, Developmental Studies Hybridoma Bank.
